# *Enterococcus faecalis* autolysin, EpaU, binds the enterococcal polysaccharide antigen via its teichoic acid-like repeats and modulates c-di-AMP signaling

**DOI:** 10.64898/2026.04.24.720668

**Authors:** Catherine T. Chaton, Nicholas R. Murner, Svetlana Zamakhaeva, Jeffrey S. Rush, Cameron W. Kenner, Alexander E. Yarawsky, Lei Huang, Parastoo Azadi, Andrew B. Herr, Natalia Korotkova, Konstantin V. Korotkov

## Abstract

The cell wall of the Gram-positive bacterium *Enterococcus faecalis* is decorated with the enterococcal polysaccharide antigen (EPA), consisting of a core rhamnan backbone linked covalently with a strain-variable teichoic acid-like (TA) polymer. Current models propose that the TA decoration is a repeating polymer composed of two alternating subunits, designated TAI and TAII, which are attached to the rhamnan core via a mild-acid labile phosphodiester bond from the initiating TAI subunit. In this study, we characterize the EpaU autolysin encoded within the EPA biosynthetic gene cluster. We demonstrate that the cell wall-binding domain of EpaU associates with the intact TA domains of EPA synthesized with the aid of the glycosyltransferases EpaR and EpaX. We further show that EpaU is a potent autolysin that binds generally over the *E. faecalis* cell surface, suggesting that it functions as a remodeling peptidoglycan hydrolase. The absence of EpaU leads to increased ampicillin resistance and elevated intracellular levels of the second messenger c-di-AMP. These data suggest that *E. faecalis* possesses a mechanism that senses the integrity of the peptidoglycan meshwork and employs c-di-AMP to regulate cell turgor, potentially altering the antibiotic resistance.

**Importance:** *Enterococcus faecalis* is an important opportunistic pathogen that can cause severe nosocomial infections. Knowledge of how bacteria remodel the cell wall is key to understanding many important cellular processes, such as antibiotic resistance, cell division, biofilm formation, and stress resistance. In this study, we shed new light on the structural details of the main cell wall polysaccharide, Enterococcal Polysaccharide Antigen (EPA), and its interaction with EpaU, an autolysin that cleaves peptidoglycan during cell wall remodeling. We also report a link between EpaU and the regulation of turgor pressure via cyclic dinucleotide signaling. This work contributes to a more complete picture of *E. faecalis* cell wall and may provide insight into the development of antimicrobial agents based on autolysins.

## Introduction

The cytoplasmic membrane of most bacteria is surrounded by a unique mesh-like polymer known as peptidoglycan, which provides structural support and shape and protects the cell from lysis caused by high internal turgor pressure. Peptidoglycan consists of polymeric strands of alternating N-acetyl glucosamine (GlcNAc) and N-acetyl muramic acid (MurNAc) disaccharides, crosslinked with short peptide chains present on MurNAc. Peptidoglycan is synthesized by multi-protein complexes consisting of penicillin-binding proteins (PBPs) and SEDS-family (Shape, Elongation, Division, and Sporulation) proteins (1, 2). To enable cell elongation and division, peptidoglycan synthesis is tightly coupled with cell wall remodeling. This continuous process involves the expansion of the peptidoglycan layer by cleaving existing peptidoglycan to create space for the insertion of new units, septal peptidoglycan splitting to separate daughter cells during division, and the release of old peptidoglycan fragments for recycling during cell wall turnover. An imbalance between peptidoglycan synthesis and cleavage can cause severe defects in bacterial cell morphology and compromise cell wall integrity, resulting in cell lysis. This mechanism underpins the action of β-lactam antibiotics, which inhibit PBPs, preventing peptidoglycan cross-linking and triggering cell lysis (3).

In Gram-positive bacteria, the secondary messenger c-di-AMP, a rheostat for turgor pressure, regulates cell wall homeostasis and resistance to β-lactam antibiotics through poorly understood mechanisms (4–11).

Bacteria commonly encode multiple secreted peptidoglycan hydrolases, also called autolysins, that participate in peptidoglycan remodeling. These enzymes are also involved in autolysis (a cell’s self-destruction) and fratricidal lysis (destruction of neighboring cells) (12–14). The last two mechanisms play important roles in the release of toxic substances, nutrients, and DNA for genetic exchange (14). Most autolysins cleave the glycosidic bond between sugar residues in glycan strands, or the glycan-peptide linkage, or the peptide-peptide linkage (14, 15). Although peptidoglycan hydrolases are significant contributors to mechanisms of resistance and tolerance to cell wall-targeting antibiotics (10, 16–18), their exact roles in these mechanisms are not fully understood due to their redundancy, organism-specificity, and complex regulation involving various cellular signals, feedback mechanisms, and environmental cues.

To prevent catastrophic cell lysis and ensure proper growth and daughter-cell separation, peptidoglycan cleavage is tightly regulated. Precise control of peptidoglycan hydrolysis is achieved by targeting autolysins to specific locations on the bacterial cell wall (12, 14). In Gram-positive bacteria, including *Enterococcus faecalis*, the cell wall contains a thick multilayered peptidoglycan decorated with capsular polysaccharides and anionic polysaccharides, such as wall teichoic acid (WTA) (19). Anionic glycopolymers play a key role in regulating peptidoglycan remodeling by serving as receptors for autolysins and/or by restricting autolysin access to certain regions of the cell wall (20–22). *E. faecalis* primarily resides in the gastrointestinal tract of animals as part of the normal gut flora (23). This organism exhibits intrinsic resistance to a wide range of cell wall-targeting antibiotics and disinfectants and can cause life-threatening infections during antibiotic-induced dysbiosis (24). *E. faecalis* peptidoglycan is decorated with Enterococcal Polysaccharide Antigen (EPA) and capsular polysaccharides (25–28). EPA plays an important role in biofilm formation, adherence to the intestinal mucus, virulence in various infection models, phage infection, and resistance to vancomycin, daptomycin, lysozyme, and other antimicrobials (29–39). Structural analysis of EPA purified from the *E. faecalis* V583 strain revealed that the polymer is composed of a polyrhamnose core which serves as a scaffold for teichoic acid-like (TA) decorations consisting of α/β-glucose (Glc), β-N-acetylgalactosamine (GalNAc), α-rhamnose (Rha), and ribitol-phosphate groups (40). The exact enzymatic steps involved in EPA initiation, elongation, transport, and assembly are not fully understood, and the current model for polymer biosynthesis is based primarily on chemical composition, NMR, and bioinformatic analysis of the EPA biosynthetic gene cluster. Most genes involved in EPA biosynthesis are present in a single locus, which is subdivided into two regions. Region 1 is composed of 18 genes (from *epaA to epaR*).

This region is highly conserved among *E. faecalis* isolates and encodes the polyrhamnose biosynthesis pathway. Region 2 is more variable and encodes the TA synthesis pathway (33, 38, 40–43). The high degree of structural diversity in EPA produced by enterococcal strains is linked to the sequence variability of Region 2 (17, 38, 40, 44). In a proposed EPA biosynthetic model, the EPA building blocks, polyrhamnose and TA decorations, are produced separately in the cytoplasm on a carrier lipid, undecaprenyl phosphate (Und-P) (40). After building blocks have been transported to the outer leaflet of the membrane and assembled into a single polymer, EPA is transferred to peptidoglycan.

Intriguingly, the TA biosynthetic gene cluster encodes the peptidoglycan hydrolase EpaU (also referred to elsewhere as AtlE (45)). A recent study reported that EpaU is an N-acetylmuramidase, which, together with the N-acetylglucosaminidase AtlA, participates in cell division septum cleavage during the stationary growth phase (46). AtlA has been reported to play a major role in cell separation during the exponential growth phase (47). The enzyme localizes specifically to the septum, splitting the peptidoglycan during cell division (48). In this study, we demonstrate that EpaU binds EPA via the TA decorations targeting specific peptidoglycan regions for cleavage. Our results provide new insights into the consequences of EpaU dysregulation, connecting peptidoglycan hydrolysis to c-di-AMP signaling and resistance to β-lactam antibiotics.

## Results

### EpaU^V583^ is a potent autolysin present as two isoforms in *E. faecalis*

*E. faecalis* EpaU consists of an N-terminal, family GH25, catalytic domain (PF01183) and a C-terminal putative cell wall binding (CWB) domain that varies greatly among *E. faecalis* strains (Fig. 1A). The N-terminal segment preceding the catalytic domain is predicted to be highly disordered (Fig. S1), with an abundance (32%) of serine and threonine residues. To understand the function of EpaU, we constructed the deletion mutant, Δ*epaU,* in the *E. faecalis* V583 variant VE14089 cured of its plasmids (49) (referred to hereafter as wild-type, WT). V583 is a clinical strain commonly used to study antibiotic resistance and virulence mechanisms. The nisin-inducible, a low-to-medium copy number, *E. faecalis* expression vector pMSP3535 (50) was employed for the complementation of Δ*epaU*. Immunoblot analysis of EpaU expression showed that incubation of WT cells with 2 % SDS effectively extracted EpaU, implying that EpaU is located on the cell surface (Fig. 2A). Interestingly, the cell-associated EpaU is present as two isoforms: a 100 kDa form, which matches the predicted molecular weight of EpaU, and a 130 kDa form. Since the N-terminal domain of EpaU is a serine/threonine-rich disordered region, it is possible that this domain is *O*-glycosylated, and the 130 kDa isoform corresponds to a glycosylated isoform of EpaU, as has been recently reported in streptococci (51). Small fragments of EpaU (70 and 55 kDa) were also found in the cell-free supernatant of WT, suggesting EpaU is actively degraded (Fig. S2A). As expected, EpaU was not detected in the Δ*epaU* supernatant (Fig. S2A) and SDS-wash sample (Fig. 2A & S2B), but was recovered in the complemented strain, Δ*epaU*::p*epaU*, upon induction with nisin in a dose-response manner (Fig. 2A, S2A, & S2B). EpaU was not induced by nisin in Δ*epaU* complemented with the empty vector (Δ*epaU*::pMSP3535) (Fig. S2B).

**Figure 1.**
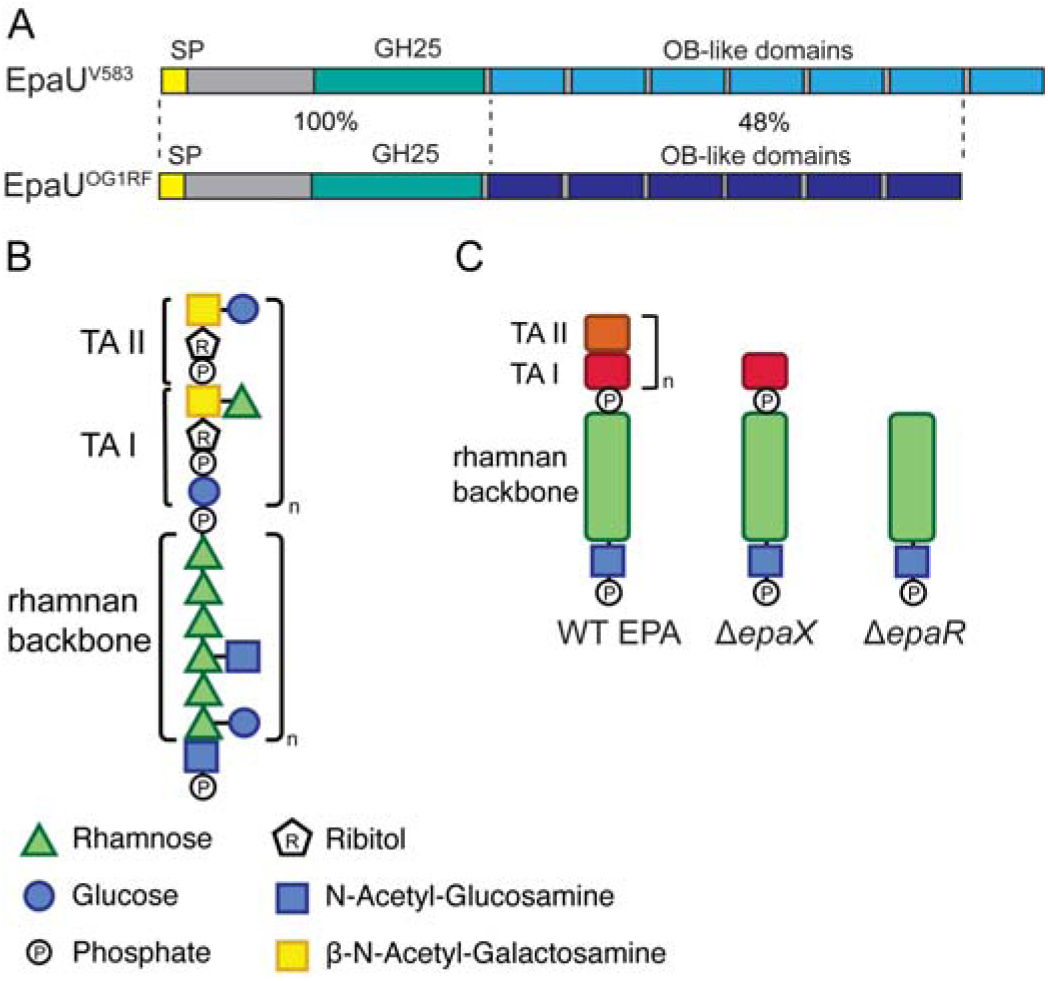
EpaU autolysin and EPA structures from *E. faecalis*. (**A**) Domain diagrams of EpaU from V583 and OG1RF strains. The percentages indicate the sequence identity between the N-terminal and the C-terminal domains of EpaU homologs from these two strains. (**B**) Previously proposed structure of EPA in *E. faecalis* V583 (40). The model proposes that EPA consists of two polymeric structures: a repeating teichoic acid-like polymer composed of alternating subunits, TAI and TAII, originating from two separate Und-P-linked lipid intermediates, and a polyrhamnnose polymer consisting of a variable number of hexa-rhamnose repeats with an unidentified number of glucose and N-acetylglucosamine side chains. The biosynthetic model is that the repeating units of the teichoic acid-like polymer are assembled on the cytoplasmic side of the plasma membrane anchored to an Und-P-P lipid anchor. After transverse diffusion to the exoplasmic surface, the repeat units are polymerized into a polymer by displacement of the phosphate molecules located at the reducing end of the incoming donor subunit by the acceptor lipid-anchored polymer, such that only the initiating subunit retains its mild acid-sensitive Glc-1-P molecule. (**C**) Simplified hypothetical model illustrating the organization of the TAI and TAII repeats in EPA derived from WT *E. faecalis* and mutant strains described in this study.

**Figure 2.**
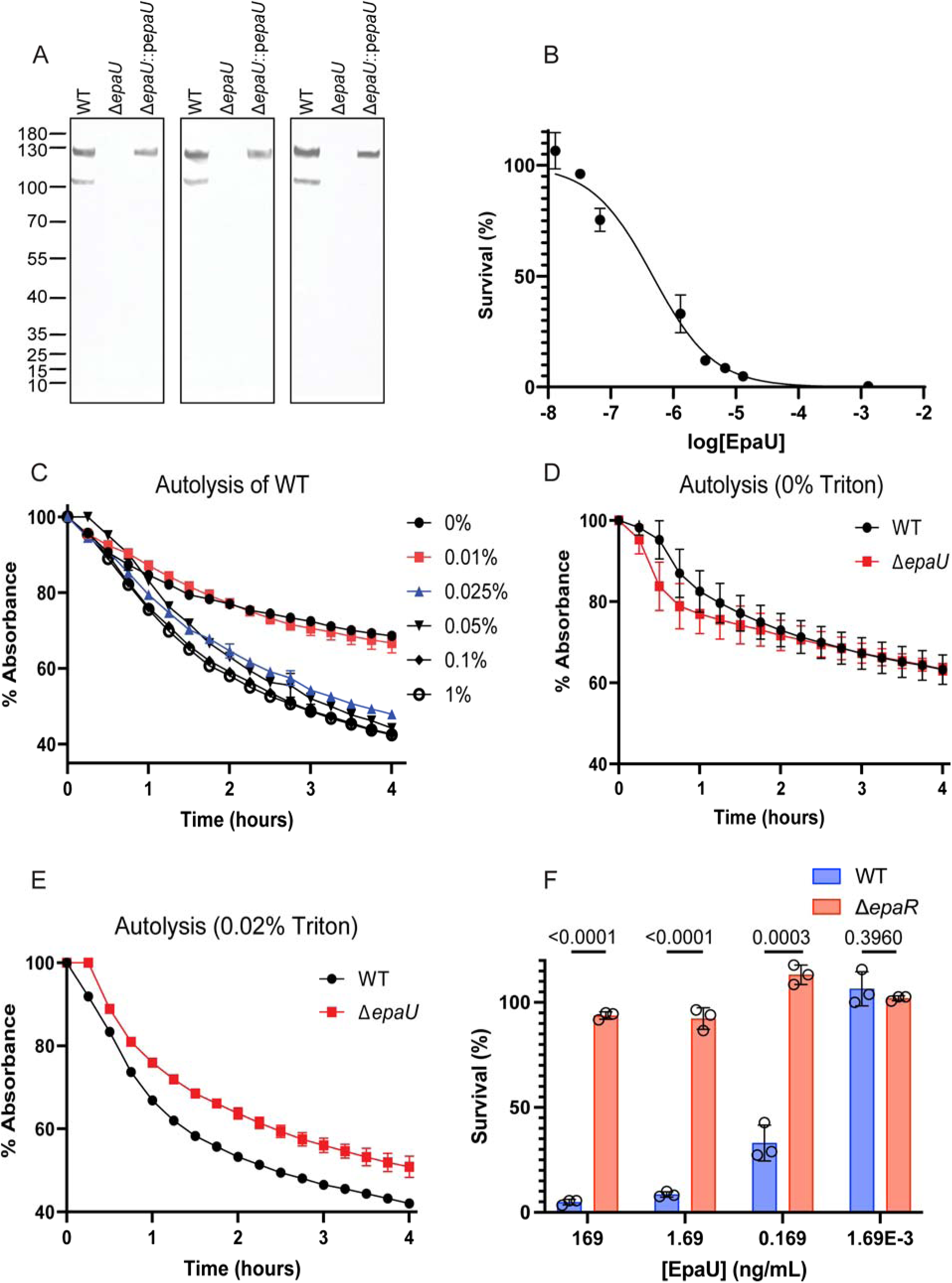
Analysis of EpaU expression in *E. faecalis* strains and activity. (**A**) Immunoblots of EpaU extracted from the cell surface of *E. faecalis* strains with 2% SDS. Proteins were separated on a 10% SDS-PAGE gel. EpaU was detected by immunoblotting using anti-EpaU antibodies. 3 independent repeats are shown. (**B**) Survival of WT bacteria with increasing concentrations of exogenous EpaU. WT Bacteria were challenged for 2 h with varying concentrations of EpaU, followed by plating for enumeration of CFUs. Error bars represent the mean and S.D., respectively (n = 3). (**C**) Autolysis of WT bacteria with varying concentrations of Triton X-100. (**D**,**E**) Autolysis of WT vs Δ*epaU* strain with 0% or 0.02% Triton X-100, respectively. For 0% Triton X-100, 3 biological repeats are represented, with error bars denoting standard error between technical repeats. For 0.02% Triton X-100, one representative repeat is shown. All biological repeats from this experiment are shown in Fig. S4, due to variation between repeats. (**F**) Bactericidal assay with exogenous EpaU. *E. faecalis* WT and Δ*epaR* were incubated with varying concentrations of EpaU for 2 h at 37 °C and then plated for enumeration of CFUs. Listed *P* values are from multiple unpaired t-tests. Columns and error bars represent the mean and S.D., respectively (n = 3).

Examination of WT and Δ*epaU* using scanning electron microscopy revealed no alterations in the cell shape of the mutant (Fig. S3), indicating that EpaU has no or a minor role in cell division. To investigate the potential function of EpaU in *E. faecalis* autolysis, we first confirmed the lytic activity of EpaU. Recombinant EpaU^V583^ purified from *E. coli* was incubated with WT *E. faecalis* cells. Bacterial survival was monitored by plating cell aliquots and counting colony-forming units (CFU) over the course of 2 h (Fig. 2B). Cell survival declined in a concentration-dependent manner, with an EC_50_ of 0.47 ng/mL, demonstrating that EpaU efficiently lyses this strain. Next, we examined whether EpaU expression in *E. faecalis* causes bacterial autolysis. To this end, we assessed the presence of the cytoplasmic HSP60 in the whole-cell and cell culture supernatant fractions of WT, Δ*epaU*, and Δ*epaU::*p*epaU* (Fig. S2C). The protein was detected in the whole-cell extract, but not in the supernatant of the analyzed strains (Fig. S2C), indicating that EpaU expression during normal growth does not trigger bacterial autolysis associated with leakage of internal contents.

Autolysis of *E. faecalis* occurs naturally, mediated by endogenous autolysins (52). Triton X-100, a membrane-permeabilizing agent that induces autolysis in Gram-positive bacteria by disrupting cell membranes, resulting in activation of autolysins associated with peptidoglycan remodeling. To understand the biological role of EpaU, we compared autolysis rates of WT and Δ*epaU* in the presence/absence of Triton X-100. Autolysis was estimated by measuring the reduction in absorbance at 600 nm (OD_600_). Incubation with Triton X-100 (0.02-1%) caused significant, time-dependent autolysis in WT (Fig. 2C). Autolysis of WT and Δ*epaU* in the absence of Triton was similar (Fig. 2D). However, compared to WT, Δ*epaU* demonstrated reduced autolysis when incubated with 0.02%, 0.2% or 1% Triton X-100 (Fig. 2E, Fig. S4).

### EpaU CWB domain recognizes TA decorations

Recent studies report that EpaU from the OG1RF strain cleaves peptidoglycan in a strain-specific manner and is inactive in an OG1RF mutant lacking the TA decorations, suggesting a role for the TA repeats in EpaU function (46). *E. faecalis* V583 and OG1RF produce structurally different EPA polysaccharides (40, 44), and the CWB domains of EpaU from these strains possess limited (48%) amino acid homology (Fig. 1A). To examine the role of the TA decorations in the lytic activity of EpaU, we constructed *E. faecalis* mutants producing EPA polysaccharides containing only the rhamnan backbone by deleting the glycosyltransferase-encoding genes *epaR* and *epaX* in the V583 variant VE14089, creating the Δ*epaR* and Δ*epaX* mutants. These genes were selected for knockout studies based on published observations that EPA variants produced by *epaR* and *epaX* mutants lack TA decorations (40, 44). EpaR has been proposed to initiate the assembly of TA decorations by transferring a glycosyl-phosphate to Und-P, thereby synthesizing the first lipid-linked intermediate in the TA biosynthetic pathway (44). EpaX has been proposed to catalyze the addition of GalNAc to the lipid-linked intermediate initiated by EpaR and to another membrane-anchored intermediate used to donate ‘building block’ units for the elongation of the TA decorations (40). When WT and the Δ*epaR* mutant were incubated with recombinant EpaU, the Δ*epaR* strain was found to be completely resistant to EpaU-mediated lysis (Fig. 2F), supporting the hypothesis that the TA decorations are critical for EpaU binding to the *E. faecalis* cell wall.

We hypothesized that the EpaU CWB domain recognizes TA variable repeats on EPA to guide enzyme localization and peptidoglycan cleavage. To test this hypothesis, we generated fluorescent fusion proteins, GFP-EpaU^V583^ and GFP-EpaU^OG1RF^, in which the CWB domains of EpaU from *E. faecalis* V583 and OG1RF strains were fused with green fluorescent protein (GFP) at the N-terminus (Fig. 3A). We used GFP-EpaU^V583^ in a co-sedimentation assay with sacculi purified from WT (V583), Δ*epaR*, Δ*epaX*, Δ*epaR*::p*epaR*, Δ*epaX*::p*epaX*, and *E. faecalis* strains — OG1RF, YI6-1, and ATCC 19433 — each with distinct Region 2 gene loci and variable CWB domains in EpaU (Fig. S5). To investigate whether the CWB domain binds *E. faecalis* capsule, the assay also included sacculi from the capsule-deficient *E. faecalis* V583 mutant, Δ*cpsC* (53). Additionally, to evaluate interactions between the CWB domain and peptidoglycan, sacculi from the *S. mutans* SCC-deficient mutant Δ*rgpG*, which produces “naked” peptidoglycan, were used (21). The strongest binding of GFP-EpaU ^V583^ was observed with WT (V583) and Δ*cpsC* (Fig. 3B). In contrast, no co-sedimentation of this fusion protein was observed with Δ*epaR*, Δ*rgpG*, OG1RF, ATCC 19433, X98, or YI6-1. Complementing Δ*epaR* and Δ*epaX* with plasmid-expressed WT copies of *epaR* and *epaX*, respectively, restored binding. These data suggest that the CWB domain of EpaU ^V583^ recognizes the V583-type TA decorations on the cell wall. To our surprise, as discussed in detail below, Δ*epaX* associated weakly with this fusion protein.

**Figure 3.**
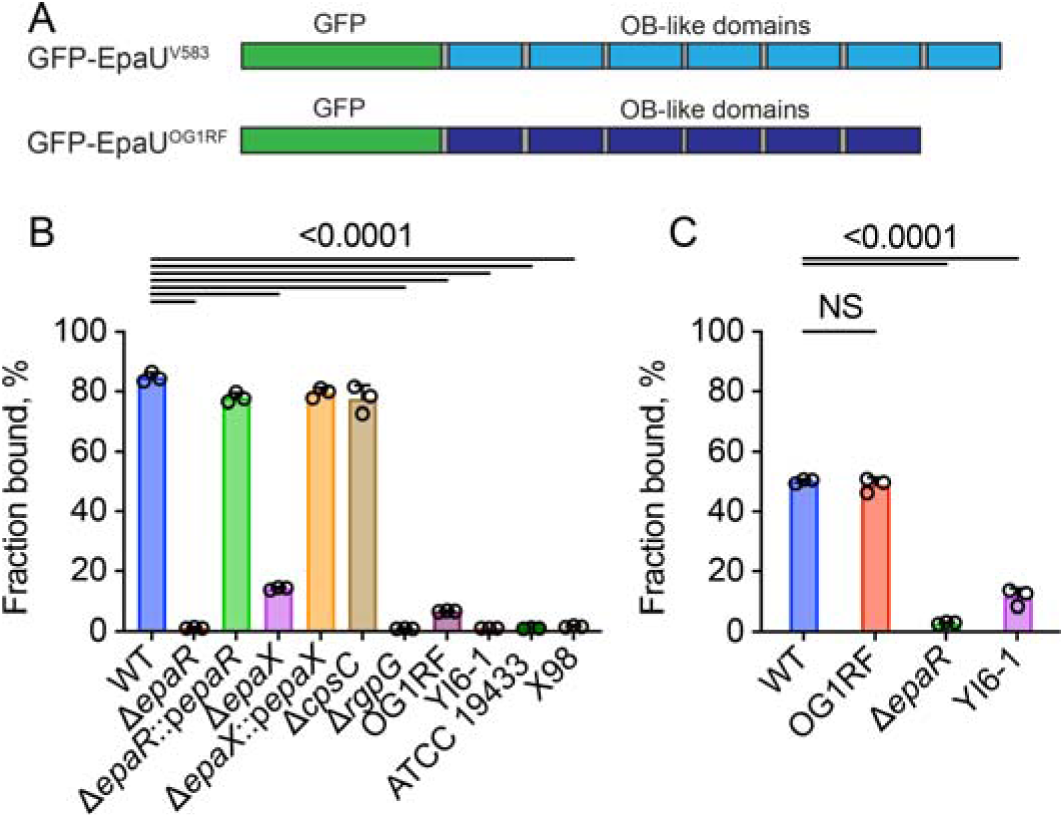
Binding of GFP-EpaU constructs to saccules from different strains. (**A**) Domain diagrams of fluorescent fusion proteins that were created by replacing the catalytic domains of EpaU with GFP. (**B**) Co-sedimentation assay of GFP-EpaU^V583^ with sacculi purified from *E. faecalis* strains. (**C**) Co-sedimentation assay of GFP-EpaU^OG1RF^ with sacculi purified from *E. faecalis* strains. Listed *P* values are from one-way ANOVA with Dunnett’s multiple comparisons test. Columns and error bars represent the mean and S.D., respectively (n = 3 in **B** and **C**).

To investigate the specificity of the CWB domain of EpaU^OG1RF^, we tested GFP-EpaU^OG1RF^ binding to sacculi purified from V583, OG1RF, Δ*epaR*, and YI6-1. As expected, this fusion protein co-sedimented with OG1RF but not with Δ*epaR*. A significant decrease in co-sedimentation of GFP-EpaU^OG1RF^ was also observed with YI6-1 sacculi. Interestingly, the association of GFP-EpaU^OG1RF^ with *E. faecalis* OG1RF and WT (V583) was not significantly different (Fig. 3C). This analysis indicates that both the CWB domains of EpaU^V583^ and EpaU^OG1RF^ recognize TA decorations, with EpaU^V583^ showing high selectivity for V583-type TA.

### EpaR and EpaX variably affect the localization of the CWB domain of EpaU to the surface of the *E. faecalis* cell

To provide a detailed picture of the bacterial regions targeted by GFP-EpaU^V583^, we incubated increasing concentrations of the fusion protein, ranging from 1.56 to 156 μg/ml, with mid-log phase WT, Δ*epaR*, Δ*epaX*, Δ*epaR*::p*epaR*, and Δ*epaX*::p*epaX* cells. The localization of GFP-EpaU^V583^ on the bacterial surface was examined using fluorescence microscopy (Fig. 4 and Fig. S6). The WT cells displayed the characteristic oval shape, with GFP-EpaU^V583^ evenly distributed across the cell surface. The Δ*epaR* cells were shorter and more spherical, suggesting a possible defect in cell division.

**Figure 4.**
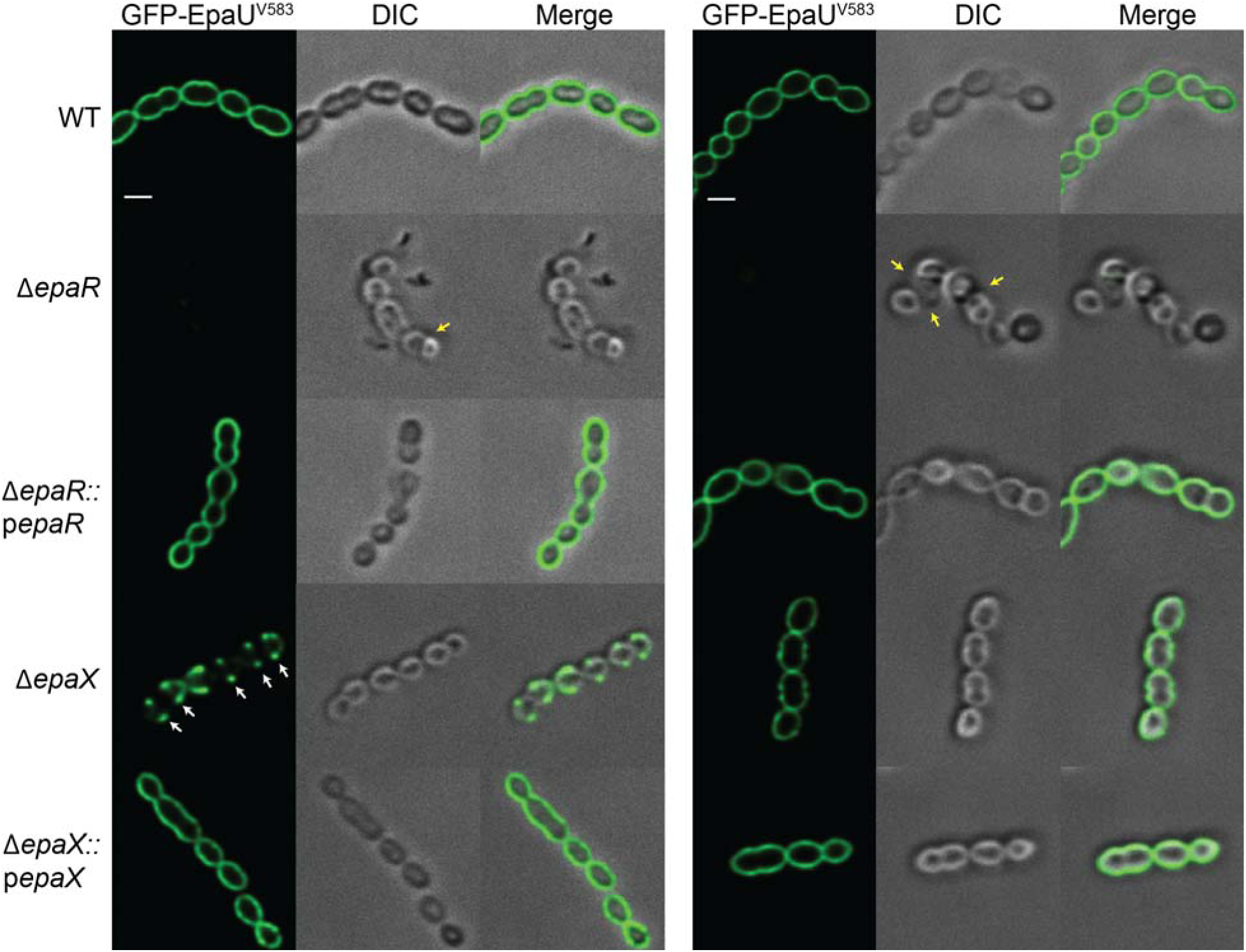
GFP-EpaU^V583^ binds differently to the cell surface of *E. faecalis* WT and mutant strains. Exponentially grown WT, Δ*epaR*, Δ*epaR*::p*epaR,* Δ*epaX* and Δ*epaX*::p*epaX* strains were incubated with GFP-EpaU^V583^ at final concentration of 1.56 µg mL^-1^ (left) or 156 µg mL^-1^ (right) for 15 min and then examined with differential interference contrast (DIC) and fluorescent microscopy. White arrows indicate equatorial sites labeled by GFP-EpaU^V583^. Yellow arrows indicate cell division defects. The experiments were performed independently three times and yielded the same results. A representative image from one experiment is shown. Scale bar is 1 µm.

However, since no discernible cell division defect was detected in the Δ*epaU* mutant, this effect is likely unrelated to EpaU binding. Consistent with the co-sedimentation assay, we did not detect GFP-EpaU^V583^ binding to Δ*epaR*, even when the protein concentration was increased a hundred-fold (Fig. 4, right panel). In contrast, the Δ*epaX* strain showed detectable levels of GFP-EpaU^V583^ binding. Unlike WT, the fluorescent signal was mainly concentrated at the equatorial sites of Δ*epaX* (Fig. 4). Increasing GFP-EpaU^V583^ concentration by a hundred-fold resulted in the loss of the distinct surface localization pattern of the fusion protein, and the fluorescent signal spread uniformly over the cell surface of Δ*epaX*. The Δ*epaR*::p*epaR* and Δ*epaX*::p*epaX* strains both appeared similar to WT, indicating restoration of the WT phenotype. These data suggest that Δ*epaX* likely produces reduced levels of TA decorations.

### EpaR is essential for TA decorations in *E. faecalis* V583

We previously reported that mild acid treatment (0.02 N HCl, 100 °C, 20 min) of streptococcal cell wall preparations releases the rhamnose-containing polysaccharides due to the presence of GlcNAc 1-phosphate in the linkage unit to peptidoglycan. The sensitivity of sugar 1-phosphates to mild acid hydrolysis is a well-known phenomenon (54), and has been reported in studies examining the linkage of WTA to peptidoglycan (55, 56). Initial experiments showed that these reaction conditions also efficiently release EPA from *E. faecalis* V583 cell walls, implying that a sugar 1-phosphate occurs in the EPA-peptidoglycan linkage region. To investigate the biosynthetic role of EpaR in EPA synthesis, EPA was released from WT and Δ*epaR* cell walls by mild acid and subsequently reduced with sodium borohydride. The polysaccharides were then analyzed for sugar composition using a gas chromatography-mass spectrometry (GC-MS) method following methanolysis in methanolic/HCl, detailed in Table 1.

**Table 1.** Compositional and reducing end sugars analysis of EPA from WT and Δ*epaR* strains.

| Strain | TMS assay |  |  |  |  | alditol acetates assay |  |  |  |
| --- | --- | --- | --- | --- | --- | --- | --- | --- | --- |
|  | Rha<sup>a</sup> | Glc | GlcNAc | GalNAc | Rbo | GlcNAcitol | GalNAcitol | Rbo | Glc-ol |
| WT | 9.1<sup>b</sup> | 4.2 | 2.4 | 2.2 | 0.7 | 1.0 | 0.3 | 0.2 | 0.05 |
| $\Delta epaR$ | 6.9 | 2.1 | 1.5 | 0.0 | 0.0 | 1.0 | 0.0 | 0.0 | 0.04 |

EPA was isolated from the WT and $\Delta epaR$ cell walls by mild acid hydrolysis, chemically reduced with sodium borohydride, and analyzed for total and reduced sugar content as described in Methods. Analysis of total sugar content was performed as TMS-derivatives of O-methyl glycosides following methanolysis with inositol internal standard. Reducing end sugar analysis was performed as alditol acetates using mannitol as an internal standard.
<sup>a</sup>Abbreviations: Rha, rhamnose; Glc, glucose; GalNAc, N-acetyl galactosamine; GlcNAc, N-acetyl glucosamine; Rbo, ribitol; N-acetyl glucosaminitol, GlcNAcitol; N-acetyl galactosaminitol GalNAcitol, Glc-ol, glucitol.
<sup>b</sup>Data are presented as a ratio to the GlcNAcitol amount.

Compositional analysis of trimethyl-silyl (TMS) derivatives of O-methyl glycosides revealed that the WT EPA mainly consists of Rha and Glc, with smaller amounts of GlcNAc, GalNAc, and ribitol (Rbo), as anticipated for a polyrhamnose polymer with Glc and GlcNAc side-chains and TA-like decorations. Importantly, EPA recovered from the Δ*epaR* cell wall preparations contained reduced amounts of Glc and GlcNAc, and no detectable GalNAc or Rbo, consistent with a polysaccharide lacking the TA-like additions. For reducing-end analysis, the reduced polysaccharides were hydrolyzed with TFA and per-acetylated to convert the reducing-end sugars into their respective alditol acetates. The analysis showed that both WT and Δ*epaR* polysaccharides contained N-acetyl glucosaminitol (GlcNAcitol) as the primary reduced sugar. Furthermore, the WT EPA contained minor amounts of N-acetyl galactosaminitol (GalNAcitol) and only trace amounts of glucitol as additional reduced sugars (Table 1). Importantly, the Δ*epaR* EPA exhibited a significantly reduced GalNAcitol content. Thus, our composition analysis supports the proposed role of EpaR in initiating the synthesis of TA decorations, and alditol acetate analysis further suggests that GalNAc is present at the reducing end of the TA decoration. These results align with the proposed initiation of EPA rhamnose core synthesis on GlcNAc-P-P-Und, as has been reported for *Streptococcus pyogenes* cell wall polysaccharides (57). However, the detection of reducing-end GalNAc raises important questions about the biosynthetic pathway model and warrants further investigation.

The released polysaccharides were further purified by size exclusion chromatography (SEC), followed by DEAE anion-exchange chromatography (AEC), revealing separation of WT EPA into two major peaks containing neutral and negatively charged polysaccharides (Fig. 5B). When AEC fractions were analyzed by GC-MS as TMS-methyl glycosides (Table 2), Rha, GlcNAc, and Glc were identified as the most abundant sugars in the neutral polysaccharide, indicating that the polyrhamnose core region is present in the first peak. Interestingly, very low levels of Rbo and phosphate were also detected in this neutral polysaccharide. The negatively charged polysaccharides were resolved into three anionic fractions by AEC using a very shallow NaCl salt gradient. As shown in Table 2, these polysaccharide species contained varying amounts of Rha and Glc, but increasing proportions of phosphate, Rbo, and GalNAc, which correlated with their binding to the DEAE column. This indicated that the anionic polysaccharide corresponds to TA decorations. To further characterize the heterogeneity of EPA WT, the polysaccharides purified by a combination of SEC and AEC were labeled at the reducing end via reductive amination with 7-amino-1,3-naphthalenedisulfonic acid (ANDS), as previously described (21). Because ANDS introduces a negative charge to polysaccharides, this fluorescent tag allows examination of the electrophoretic mobility of both neutral and charged polysaccharides by polyacrylamide gel electrophoresis (PAGE). Consistent with the AEC analysis, ANDS-labeled TA decorations displayed the characteristic “laddering” pattern of fluorescent bands, indicating the high level of heterogeneity of the polysaccharide (Fig. 5D). As expected, the bands of the TA species containing the highest level of phosphate were the fastest migrating in these conditions. This observation suggests that the recovered TA decorations are a mixture of polysaccharides with repeat units of varying degrees of polymerization.

**Figure 5.**
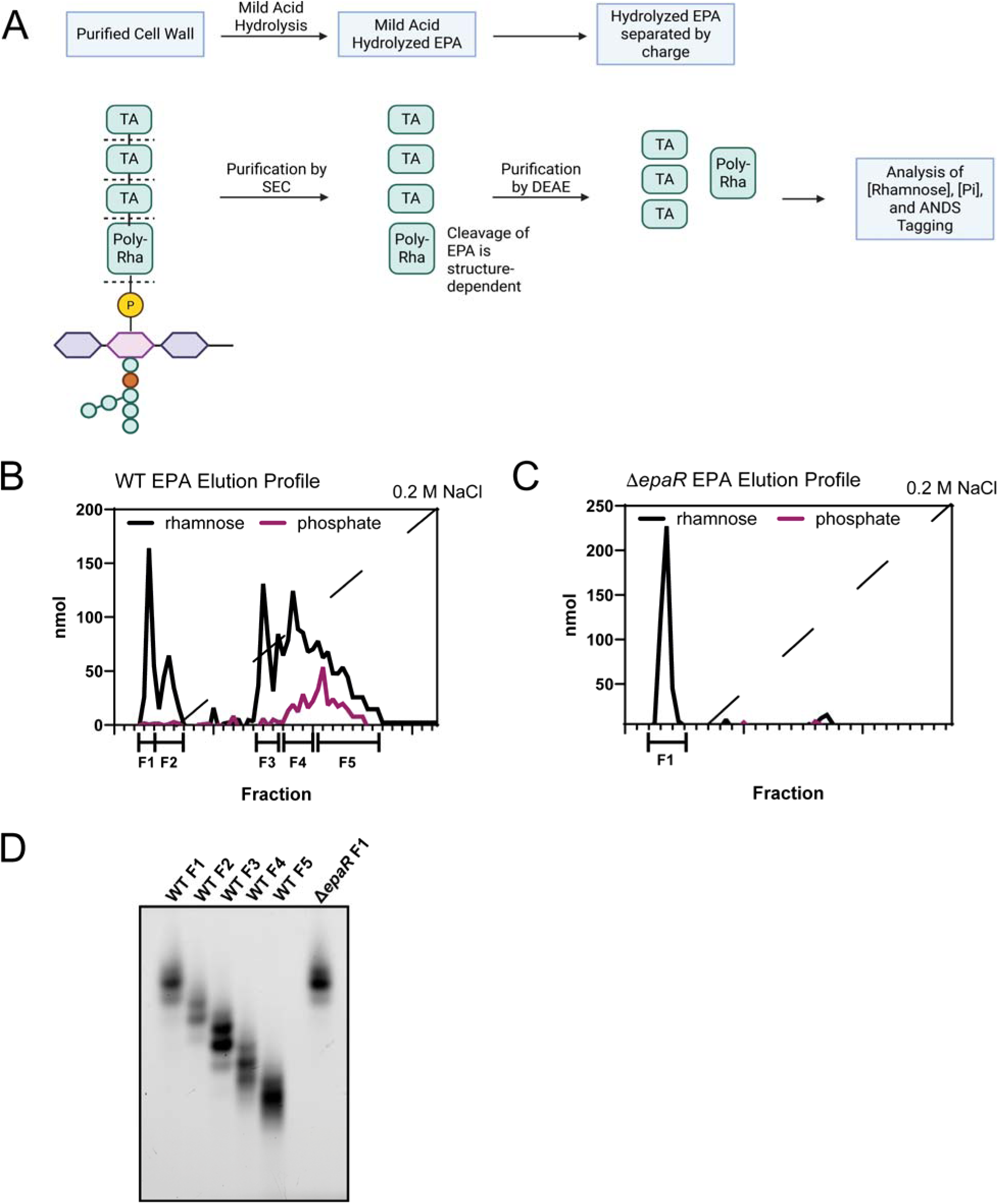
Purification and analysis of WT and Δ*epaR* EPA released from cell walls by mild acid hydrolysis. (**A**) Workflow illustrating the purification and analysis of EPA variants released from cell walls by mild acid hydrolysis (0.02 N HCl at 100 °C for 20 min). Dashed lines mark sites of hydrolysis of phosphodiester bonds that link EPA to peptidoglycan and TA decorations to a rhamnan backbone. (Created in BioRender Murner, N. (2026) https://BioRender.com/r84yrou). (**B** and **C**) EPA from WT (B) and Δ*epaR* (C) released from cell walls by mild acid was fractionated by size-exclusion chromatography (SEC) on Biogel P150, as described in Methods. Fractions containing EPA were combined, concentrated, desalted using a spin column (Amicon 3,000 MWCO filters), and resolved by anion exchange chromatography on an 18 ml column of DEAE Toyo 650M equilibrated in 5 mM HEPES-NaOH, pH 7.4. The columns were eluted with 40 mL of starting buffer and then with a gradient (80 mL) of NaCl (0-0.2 M, dashed line). Fractions of 2 mL were collected and analyzed for carbohydrate (solid black line, anthrone analysis) and phosphate (solid purple line, malachite green analysis) as described in Methods. (**D**) DEAE fractions were combined as indicated, fluorescently-tagged at the reducing end by reductive amination with ANDS, resolved by SDS-PAGE, and visualized by fluorescent imaging analysis. Individual species are resolved based on the electromotive force contributed by the presence of varying numbers of the phosphate-containing TA repeat units. In **B, C** and **D**, the experiments were performed independently two times and yielded the same results. A representative image from one experiment is shown.

**Table 2.**
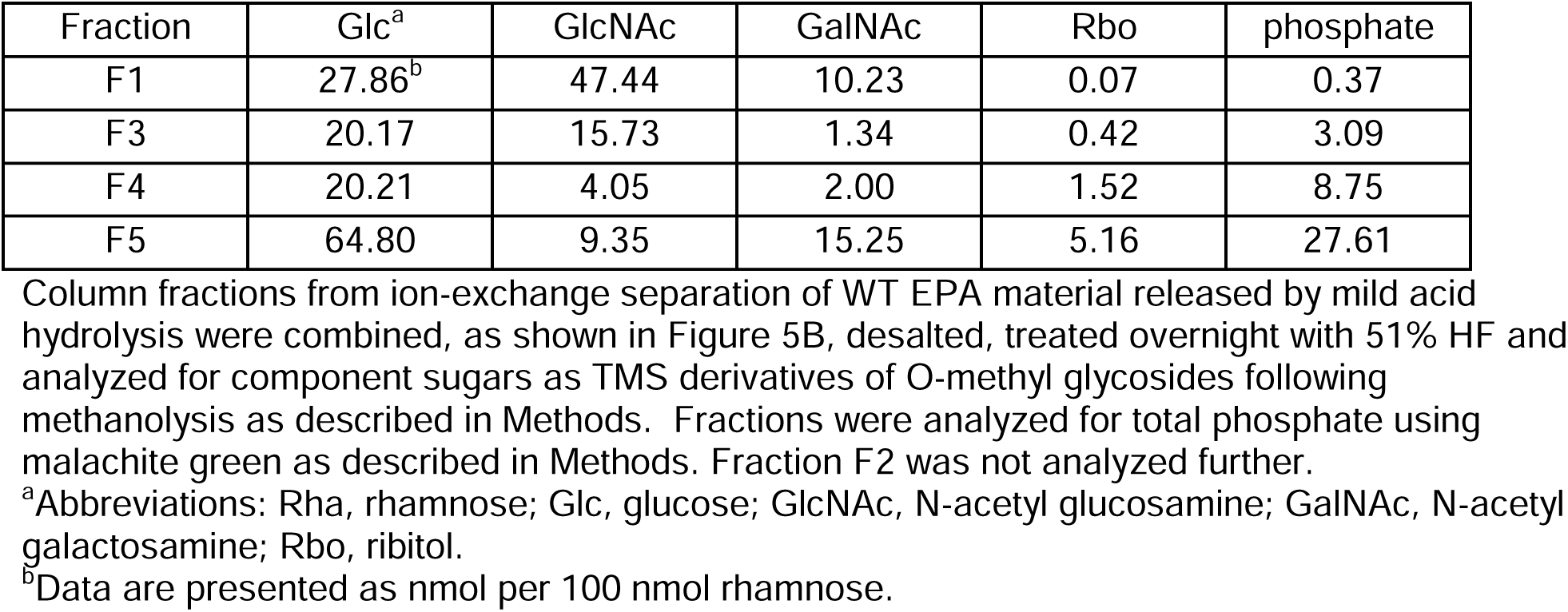
Compositional analysis of EPA fractions separated by ion-exchange chromatography.

| Fraction | Glc<sup>a</sup> | GlcNAc | GalNAc | Rbo | phosphate |
| --- | --- | --- | --- | --- | --- |
| F1 | 27.86<sup>b</sup> | 47.44 | 10.23 | 0.07 | 0.37 |
| F3 | 20.17 | 15.73 | 1.34 | 0.42 | 3.09 |
| F4 | 20.21 | 4.05 | 2.00 | 1.52 | 8.75 |
| F5 | 64.80 | 9.35 | 15.25 | 5.16 | 27.61 |

<sup>a</sup>Abbreviations: Rha, rhamnose; Glc, glucose; GlcNAc, N-acetyl glucosamine; GalNAc, N-acetyl galactosamine; Rbo, ribitol.
<sup>b</sup>Data are presented as nmol per 100 nmol rhamnose.

In contrast to WT EPA, a single peak of neutral material was detected when the EPA variant extracted from Δ*epaR* by mild acid treatment was purified using AEC (Fig. 5C). Furthermore, the electrophoretic mobility of the ANDS-labeled Δ*epaR* EPA matched that of the WT neutral polysaccharide (Fig. 5D), indicating that the WT neutral polysaccharide corresponds to the rhamnan backbone core of EPA.

### The Δ*epaX* EPA has fewer TA decorations than the WT EPA

Previously, it has been reported that the V583 EPA treated with hydrofluoric acid (HF) can be separated by SEC into a rhamnan linkage core and a low-molecular-weight polymer identified as TA decorations (40). These data indicate that TA decorations are linked to a rhamnan backbone via HF-labile phosphodiester bonds in this polysaccharide. Because the SEC elution profile of the HF-treated Δ*epaX* EPA did not reveal a low-molecular-weight component, it has been suggested that this EPA variant contains only a rhamnan backbone (40). Based on our findings that the CWB domain of EpaU^V583^ binds to Δ*epaX*, we hypothesized that the Δ*epaX* EPA contains a minor low-molecular-weight TA decoration that elutes in the low-molecular-weight inclusion volume during SEC, leading to its loss. To test this idea, we modified the EPA purification protocol to enzymatically release soluble polysaccharide fragments from cell walls using an N-acetylmuramidase, mutanolysin, and D-alanyl-L-alanine endopeptidase, zoocin A (58) (Fig. 6A). After cell wall polysaccharides (EPA and capsular material) were cleaved from WT, Δ*epaR*, and Δ*epaX* cell walls, they were subjected to composition analysis. The WT polysaccharide material contained Rha and Glc as the predominant sugars with smaller amounts of GalNAc, GlcNAc, and ribitol. Compared to WT, the Δ*epaR* and Δ*epaX* samples displayed no detectable ribitol, and reduced amounts of Glc and GalNAc (Table 3). These data indicate that both the Δ*epaR* and Δ*epaX* EPA have defects in TA decorations. We further separated EPA from capsular polysaccharides by SEC. To free EPA from peptidoglycan fragments, we treated the polysaccharide material with mild acid as described above. Finally, EPA was purified by AEC and fluorescently labeled with ANDS (Fig. 6A). We observed that the WT EPA variant was separated into a large peak containing relatively neutral polysaccharide (WT F1 fraction) and multiple peaks containing negatively charged polysaccharide (WT F2 and WT F3) (Fig. 6B). As expected, PAGE analysis revealed that ANDS-labeled WT F1 and WT F2 are heterogeneous polysaccharides of different sizes (Fig. 6E). Interestingly, WT F2 and WT F3 contained polysaccharide material running close to the dye front, suggesting the presence of a low-molecular-weight decoration cleaved off by mild acid hydrolysis. We did not detect these small-sized components when EPA released from peptidoglycan by mild acid hydrolysis was subjected first to SEC purification (Fig. 5D), suggesting their loss during this step.

**Figure 6.**
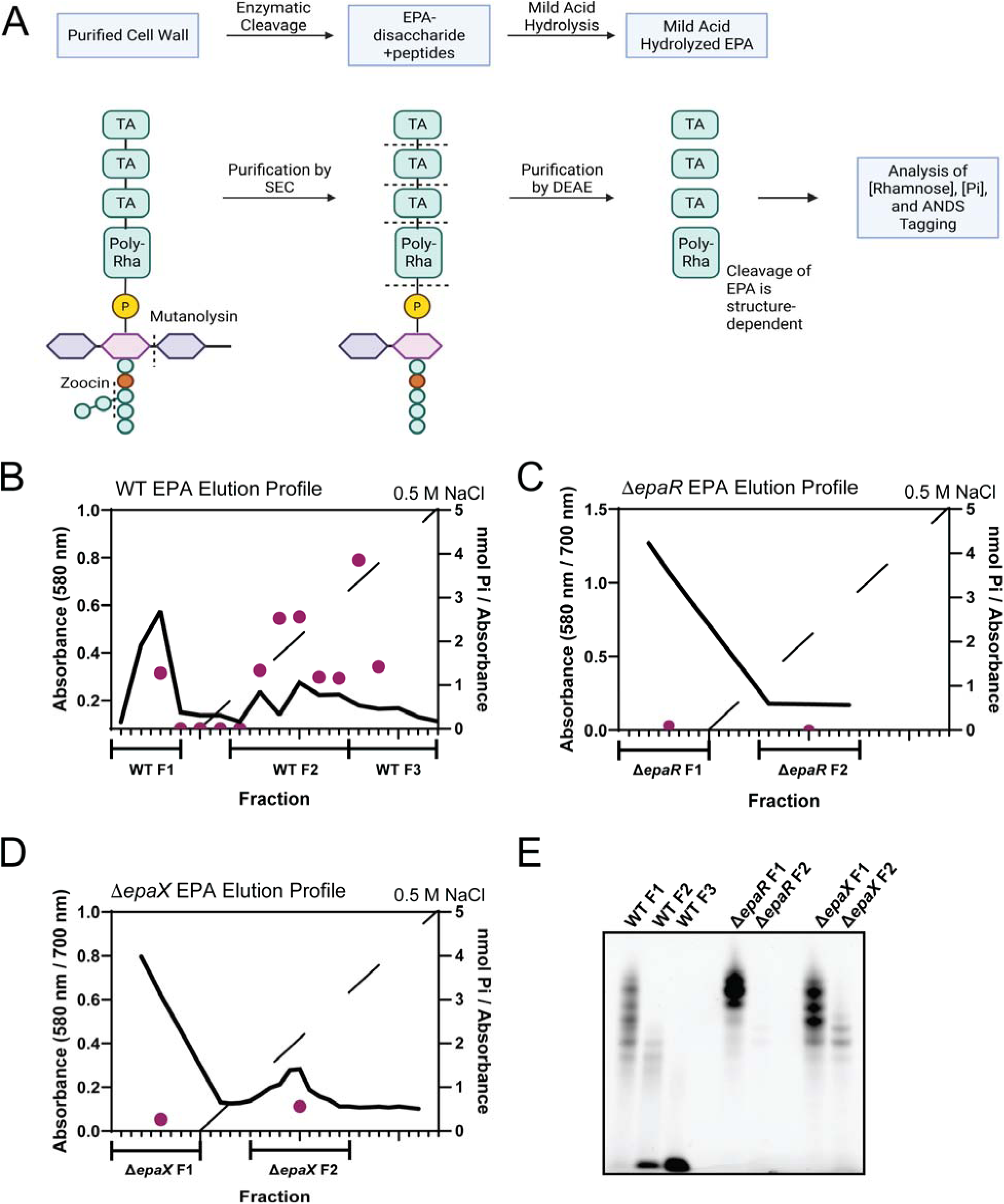
Purification and analysis of WT, Δ*epaX*, and Δ*epaR* EPA variants released from cell walls by enzymatic cleavage. (**A**) Workflow illustrating the purification and analysis of EPA variants released from cell walls by mutanolysin and zoocin A. EPA material was purified by size-exclusion chromatography (SEC), followed by mild acid hydrolysis to cleave EPA from the disaccharide-peptide fragment of peptidoglycan and TA decorations from the rhamnan backbone. Dashed lines mark sites of enzymatic cleavage and mild acid hydrolysis of phosphodiester bonds linking EPA to peptidoglycan and TA decorations to a rhamnan backbone. (Created in BioRender. Murner, N. (2026) https://BioRender.com/19xslx3). (**B-D**, left axis) The elution profiles from a DEAE chromatography column as measured by a colorimetric anthrone assay, where 580 nm (solid line) detects primarily Rha, 700 nm (dashed line) detects primarily Glc (**B-D**, right axis), and the diagonal dashed line indicates the salt gradient. Phosphate assay (purple dots) of select fractions of each elution profile. (**E**) ANDS labeled SDS-PAGE gel of pooled fractions. After mild acid hydrolysis, a different number of species are formed depending on the amount of phosphate-containing TA repeats present in EPA, with WT having a significant number, Δ*epaR* essentially having almost no TA decoration, and Δ*epaX* having a significantly reduced number. In **B, C, D** and **E**, the experiments were performed independently two times and yielded the same results. A representative image from one experiment is shown.

**Table 3.** Compositional analysis of EPA from WT and mutant strains.

| Strain | Rha<sup>a</sup> | Gal | Glc | GalNAc | GlcNAc | Rbo |
| --- | --- | --- | --- | --- | --- | --- |
| WT | 100.0<sup>b</sup> | 0.3 | 39.4 | 16.4 | 7.2 | 0.4 |
| $\Delta epaX$ | 100.0 | 0.1 | 13.9 | 4.4 | 10.4 | ND<sup>c</sup> |
| $\Delta epaR$ | 100.0 | 0.1 | 10.5 | 3.3 | 13.0 | ND |

EPA was released from the WT, $\Delta epaR$ and $\Delta epaX$ cell walls by incubation with mutanolysin and zoocin A, as described in Methods. Glycosyl composition was performed by the CCRC as TMS-derivatives of O-methyl glycosides following methanolysis with inositol internal standard.
<sup>a</sup>Abbreviations: Rha, rhamnose; Gal, galactose; Glc, glucose; GalNAc, N-acetyl galactosamine; GlcNAc, N-acetyl glucosamine; Rbo, ribitol.
<sup>b</sup>Data are presented as pmol per 100 pmol of Rha.
<sup>c</sup>ND, not detected.

Both the Δ*epaR* and Δ*epaX* EPA variants, purified using the modified protocol (Fig. 6C, D), contained the large peak of relatively neutral polysaccharide corresponding to the WT F1 fraction (Fig. 6E). However, Δ*epaR* EPA had very little carbohydrate or phosphate in the later fractions (Fig. 6C), while the Δ*epaX* EPA (Fig. 6D) had a single, smaller retained peak compared to WT. Furthermore, the electrophoretic mobility of the ANDS-labeled Δ*epaR* F1 and Δ*epaR* F2 fractions revealed a major band with relatively few released products in the first fraction and almost no signal in the second fraction.

Importantly, PAGE analysis of the ANDS-labeled Δ*epaX* F1 and Δ*epaX* F2 fractions demonstrated “laddering” of the polysaccharide similar to WT F1 and WT F2, indicating that this variant is decorated with negatively charged moieties (Fig. 6E). However, a low-molecular-weight polysaccharide fragment was absent in Δ*epaX* F2 (Fig. 6D, E), suggesting that EpaX participates in the synthesis of this decoration. Taken together, these observations support the idea that EPA consists of two types of TA decorations: TAI, which requires EpaR activity, and TAII, which depends on EpaX activity (Fig. 1B, C).

### TA decorations modulate interactions of the EpaU CWB domain with EPA

To gather additional evidence that the CWB domain of EpaU interacts with TA decorations present on the WT and Δ*epaX* EPA variants, we investigated the complex formation between GFP-EpaU^V583^ and EPA variants by analytical ultracentrifugation employing both continuous c(s) distribution and wide distribution analyses. The EPA variants were released from the WT, Δ*epaR*, and Δ*epaX* cell walls by enzymatic digestion or mild acid treatment, and then purified using SEC. When the enzymatically digested WT EPA was mixed with GFP-EpaU^V583^, multiple high-molecular-weight species were observed between 5-13 S (Fig. 7A, B), indicating that EpaU recognizes EPA, and forms binding configurations with different stoichiometries. In contrast, the enzymatically digested Δ*epaX* EPA incubated with GFP-EpaU^V583^ revealed substantially smaller complexes sedimenting at 6.5 S (Fig. 7A, B). This observation is consistent with the characterization of Δ*epaX* EPA by AEC and electrophoretic mobility analysis outlined above, implying that Δ*epaX* EPA has fewer TA decorations, leading to a lack of higher order complexes. In the case of the enzymatically digested Δ*epaR* EPA, the primary species was observed at just below 5 S, the same location as the EpaU control, indicating no complex formation (Fig. 7A, B).

**Figure 7.**
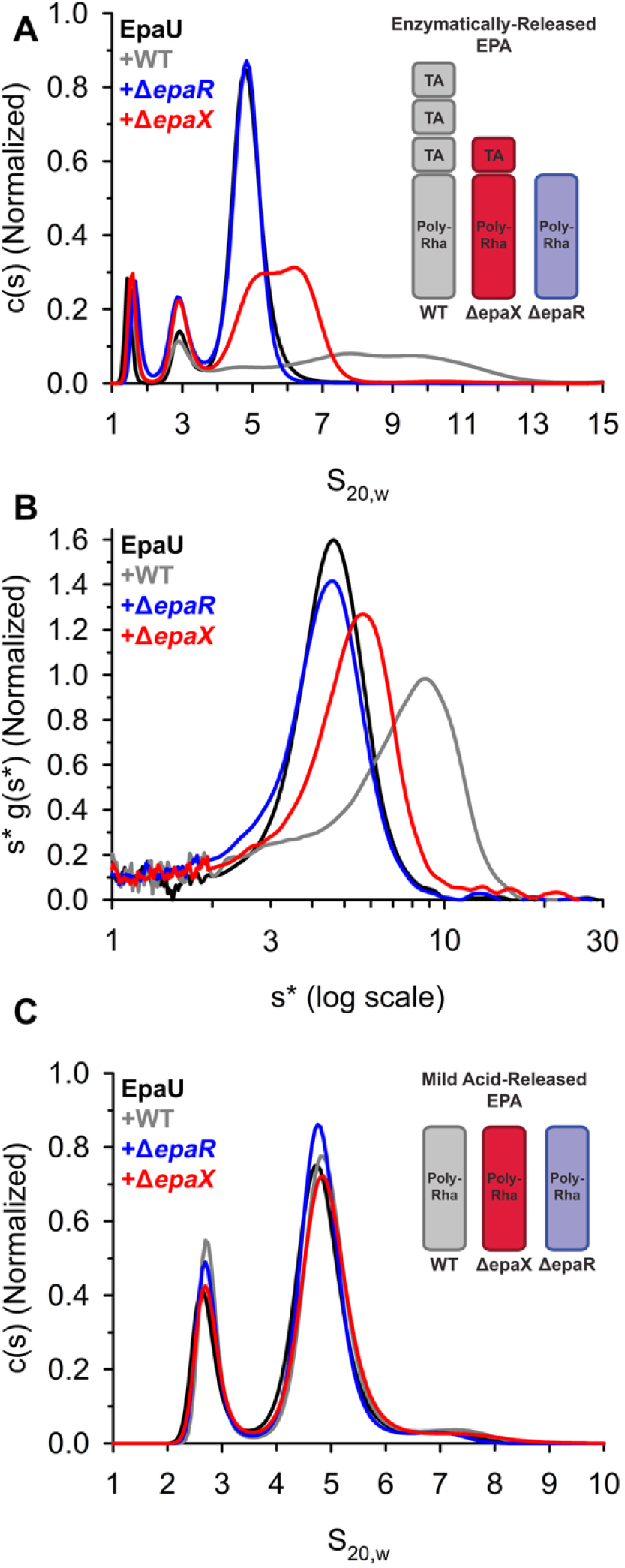
Analytical ultracentrifugation analysis of GFP-EpaU^V583^ binding to EPA variants released from cell walls of *E. faecalis* strains by different methods. (**A**) The continuous c(s) distributions and (**B**) wide distribution analysis (plotted as s*g(s*) vs. s* on a log scale) for EpaU alone (black line) and combined with EPA variants that were released from WT (grey line), Δ*epaR* (red line), and Δ*epaX* (purple line) by enzymatic cleavage. (**C**) The c(s) distribution of EpaU alone (black line) and combined with EPA variants that were released from WT (grey line), Δ*epaR* (red line), and Δ*epaX* (purple line) by mild acid hydrolysis. The experiments were performed independently two times and yielded the same results. A representative image from one experiment is shown.

Importantly, when the polysaccharides were released from the cell wall using mild acid hydrolysis, no complexation was detected (Fig. 7C). These data suggest that mild acid cleaves TA decorations from EPA, rendering the polymer unrecognizable by the CWB domain of EpaU.

### Deletion of EpaU increases resistance to ampicillin and the intracellular concentration of c-di-AMP

To understand how the inactivation of EpaU affects the susceptibility of *E. faecalis* V583 to antibiotics interfering with the synthesis of the cell wall, we measured MIC values of WT and Δ*epaU* for ampicillin, penicillin, cephalosporin, ceftriaxone, vancomycin, daptomycin, or tunicamycin (Table S1). Compared to WT, the Δ*epaU* mutant exhibited increased resistance to ampicillin (Fig. 8A, B). MIC values of WT and Δ*epaU* were 6.986 and 10.61 μg/mL, respectively (Table 4). In contrast, the ampicillin resistance of the AtlA deletion mutant was similar to that of the V583 background strain (Fig. S7). These data suggest that inactivation of EpaU strengthens the cell wall, thereby increasing bacterial resistance to ampicillin, which inhibits peptidoglycan cross-linking.

**Figure 8.**
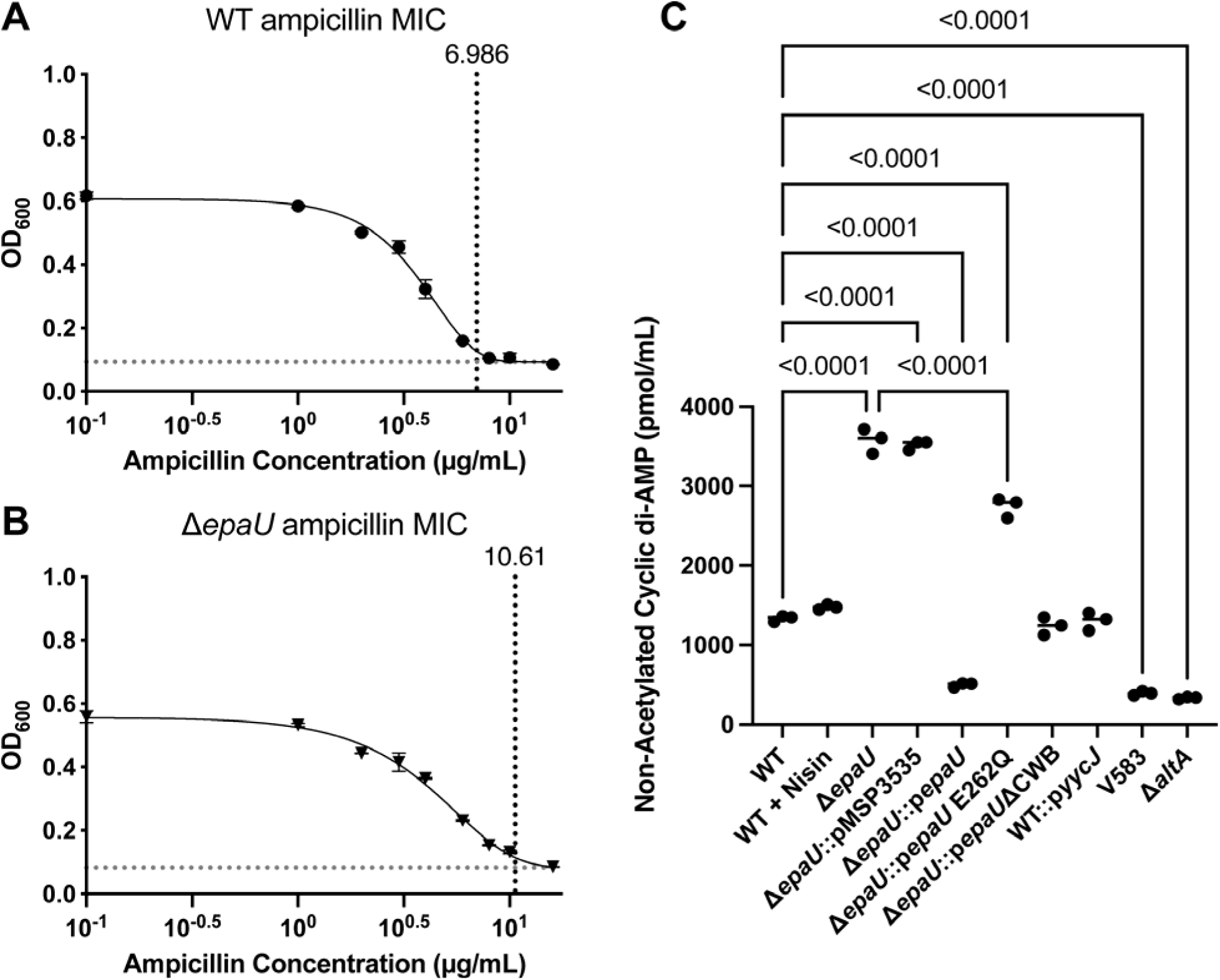
The absence of EpaU leads to increased resistance to ampicillin and elevated intracellular levels of c-di-AMP. (**A, B**) Dose-response curves for ampicillin resistance of *E. faecalis* WT (A) and the Δ*epaU* mutant (B). Bacterial growth data (OD_600_ nm) were fitted to a modified Gompertz model. The vertical dotted line shows the calculated threshold at which the growth is inhibited (MIC). Error bars represent standard deviation across three repeats. (**C**) Intracellular levels of cyclic di-AMP in the WT and Δ*epaU* cells. An empty nisin-inducible vector, pMSP3535 was used alongside both WT and Δ*epaU* backgrounds to express an unrelated YycJ enzyme as a negative control, along with a plasmid complementation of *epaU* in the Δ*epaU* background, in addition to a predicted catalytic mutant *epaU* E262Q and an *epaU* which lacks the cell-wall binding domain (*epaU* ΔCWB). The V583 strain was also measured alongside Δ*altA*. Listed *P* values are from one-way ANOVA. All experiments were done in triplicate, and the line represents the median value with WT and V583 standards replicated for each ELISA.

**Table 4.**
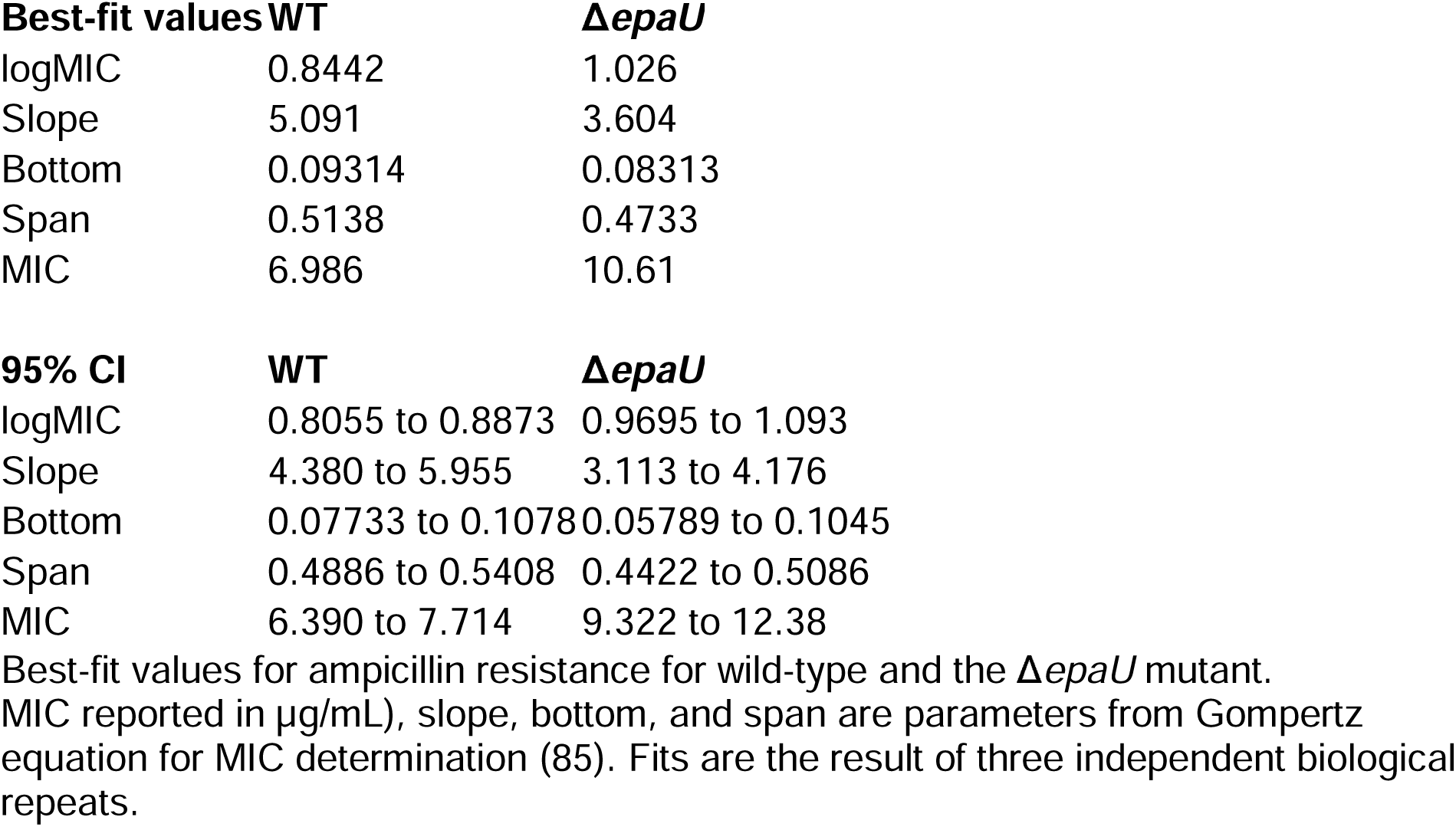
Ampicillin resistance of the WT and Δ*epaU* strains.

The signaling molecule c-di-AMP regulates cytoplasmic turgor pressure and helps maintain appropriate bacterial cell size and volume. In *E. faecalis*, it is synthesized by the diadenylate cyclase DacA encoded by *cdaA* (59). A recent study demonstrated that *B. subtilis* actively monitors its peptidoglycan meshwork and, in response to defects, activates c-di-AMP synthesis (11). We hypothesized that increased structural integrity of peptidoglycan in Δ*epaU* might be sensed by the *E. faecalis* cell, triggering a reduction in c-di-AMP levels. To test this, we measured intracellular c-di-AMP concentrations in WT, Δ*epaU*, Δ*epaU*::p*epaU*, and Δ*epaU*::pMSP3535 (Fig. 8C, S8). As a positive control, we constructed a strain WT::p*cdaA*, carrying a nisin-inducible *cdaA*. The amount of c-di-AMP detected in WT was 1330 ± 37.6 pmol/ml (n=3). When expression of DacA was induced by the addition of 25 nM nisin, c-di-AMP levels increased by approximately 3-fold. Surprisingly, deletion of EpaU resulted in a three-fold rise in c-di-AMP levels. The cellular c-di-AMP concentrations decreased in a dose-response manner when Δ*epaU*::p*epaU* strain was grown in the presence of increasing concentrations of nisin (Fig. S9), confirming that cellular c-di-AMP levels respond to EpaU expression (Fig. S2B). Supplementation of the medium with 3.12 ng/ml nisin resulted in 785 ± 21.9 pmol/ml c-di-AMP, and it reached 337 ± 16.1 pmol/ml c-di-AMP upon EpaU induction with 25 ng/ml nisin (Fig. S9). In contrast, incubation of the WT or Δ*epaU*::pMSP3535 strains with 25 ng/ml nisin did not affect their c-di-AMP levels. Interestingly, compared to the *E. faecalis* V583 background strain, the intracellular c-di-AMP concentration in Δ*atlA* was not altered (Fig. 8C). Furthermore, when we exposed WT *E. faecalis* to varying concentrations of lysozyme, no significant increase in c-di-AMP levels was observed (Fig. S10, Table S2), indicating that EpaU cleavage of peptidoglycan generates specific signals recognized by the c-di-AMP signaling mechanism which are not simulated by AtlA or lysozyme. Ampicillin MIC values for WT::p*cdaA* remained similar to those of WT (Fig. S7, Table S2), which demonstrates that increased cellular c-di-AMP levels do not confer increased ampicillin resistance in *E. faecalis*.

To investigate how EpaU catalytic activity and its binding to EPA impact c-di-AMP levels, we complemented Δ*epaU* with the catalytically inactive *epaU* (Δ*epaU*::p*epaU*-E262Q) and the *epaU* variant lacking the CWB domain (Δ*epaU*::p*epaU*ΔCWB) expressed under the nisin-inducible promoter. The c-di-AMP content in Δ*epaU*::p*epaU*-E262Q induced with 25 ng/ml nisin was slightly lower than in Δ*epaU*, but c-di-AMP concentration did not decrease to levels detected in WT cells (Fig. 8C). Interestingly, the induction of EpaUΔCWB with 25 ng/ml nisin fully restored c-di-AMP in Δ*epaU*::p*epaU*ΔCWB to levels observed in the WT strain (Fig. 8C). However, 5 ng/ml nisin did not affect the c-di-AMP content in this strain (Fig. S11). Thus, our findings demonstrate that, in response to peptidoglycan cleavage mediated by the EpaU catalytic domain, *E. faecalis* reduces the intracellular concentration of c-di-AMP, and that binding of EpaU to EPA via the CWB domain is necessary for efficient peptidoglycan cleavage, thereby producing a robust effect on c-di-AMP levels.

## Discussion

Enterococcal infections have become one of the most challenging nosocomial problems worldwide due to their intrinsic and acquired resistance to many antibiotics, including a large group of β-lactams (60–62). Insights into the functions and regulatory mechanisms of autolysins in *E. faecalis* could identify effective avenues for developing compounds that activate or facilitate bacterial lysis during infection. Although *E. faecalis* genomes encode multiple putative peptidoglycan hydrolyses (63), only a few have been characterized in detail. A recent study conducted in reference strain OG1RF demonstrated that the autolysin EpaU, encoded by a variable genetic locus within the EPA biosynthesis gene cluster, is an N-acetylmuramidase that contributes to the cleavage of peptidoglycan and plays a role in the pathogenesis of *E. faecalis* (46). EpaU possesses a highly variable CWB domain, suggesting its role in targeting the strain-specific TA decorations. In this work, we confirm that EpaU functions as an active autolysin in *E. faecalis* V583, as well. Using pull-down assays and analytical ultracentrifugation to examine the association of the fluorescently-labeled CWB domain with *E. faecalis* V583 and the TA-deficient mutants, Δ*epaR* and Δ*epaX*, we revealed that this domain binds EPA tightly, recognizing the TA decorations. The structural epitope recognized by the domain likely consists of a few intact TA repeating units because mild acid hydrolysis of EPA completely destroyed the association between CWB and EPA. When evaluating the interaction of the CWB domain with sacculi purified from different *E. faecalis* isolates, we observed that the variant from V583 is highly specific for its own cell wall compared with the variant from *E. faecalis* OG1RF, indicating strain-specific differences in the mechanism of EPA recognition by the CWB domain of EpaU.

A recent study has reported that EpaR and EpaX glycosyltransferases encoded within the core and variable gene loci of the V583 EPA biosynthesis gene cluster, respectively, are essential for TA biosynthesis (40, 44). It was then quite surprising that Δ*epaX*, which was previously reported to lack TA decorations, still exhibited binding of the protein fusion, although significantly diminished in comparison to WT cells. We observed that binding of the fluorescently-labeled CWB domain to the Δ*epaX* mutant was restricted to the equatorial rings when the protein was titrated downward. The analysis of the electrophoretic mobility of EPA variants produced by WT and the EPA mutants demonstrated that the Δ*epaX* EPA contains a reduced number of TA decorations. These results are consistent with a hypothetical model of the V583 EPA structure (40) in which the TA domain is composed of a repeating polymer assembled from two building-block subunits referred to as TAI and TAII (40). In this model, TAI synthesis is initiated by EpaR, forming Glc-P-P-Und, and EpaX is required to add GalNAc to both the TAI and TAII subunits. Alternatively, in a recent study (44), EpaR has been proposed to catalyze the synthesis of GalNAc-P-P-Und, which serves as an initiator of the TAI subunit. Our alditol acetate analysis supports the proposed function of EpaR and further suggests that the TAI subunit is likely initiated with a GalNAc residue. This second model of OG1RF EPA structure disagrees with a previously proposed model (40) in which multiple repeats of the TA decoration consist of a TAI-TAII hybrid connected to the polyrhamnose core by the Glc residue originating from the reducing end of the TAI intermediate (Fig. 1B). However, these competing models are somewhat speculative and further analysis will be required to complete the characterization of the V583 and OG1RF EPA structures. Based on our data, we propose a model of V583 EPA biosynthesis, in which GalNAc transferase EpaR initiates TAI unit assembly, which can be elongated by the addition of TAII-dependent subunits, or can be transferred to the rhamnan backbone in the absence of EpaX. EpaX is proposed to initiate the synthesis of the TAII unit, which is required for TAI-TAII repeat unit assembly, i.e. EpaX activity enables TA polymerization.

Fluorescence microscopy analysis of the association between the EpaU^V583^ CWB domain and bacterial cells revealed that the domain binds evenly across the cell surface of WT bacteria, suggesting that EPA is decorated with TA throughout the cell surface.

The distinct binding of the EpaU^V583^ CWB domain to the equatorial rings of Δ*epaX* could be explained by the observation that the newly synthesized peptidoglycan layer at cell division sites is generally thinner than in other areas of the cell wall to allow for daughter cell separation during division (64). It has been shown that, similar to the EPA structure, the rhamnose-rich polysaccharide of *Lactococcus lactis* MG1363 consists of two polysaccharides, a linear α-L-rhamnan polymer and a negatively-charged polysaccharide pellicle, covalently linked together into a single heteropolysaccharide (65, 66). The polysaccharide pellicle forms an outer layer coating the bacterial cells, and the rhamnan polymer is trapped inside the peptidoglycan layer (65). It is possible that in *E. faecalis*, EPA has a similar surface organization. If the variable TA decorations are surface-exposed, but the rhamnan core is embedded within the peptidoglycan layer, the Δ*epaX* EPA would be unmasked for EpaU binding at the division site. This idea is also consistent with the observation that the EpaU^V583^ CWB domain binds uniformly across the cell surface of the Δ*epaX* EPA when the CWB domain concentration is significantly increased.

Our results further demonstrate that the absence of EpaU hydrolase activity in *E. faecalis* reduces sensitivity to ampicillin and increases cellular c-di-AMP levels. The CWB domain of EpaU contributes to the regulation of c-di-AMP levels, underscoring the importance of EpaU binding to EPA for efficient peptidoglycan cleavage and maintenance of c-di-AMP homeostasis. In bacteria, c-di-AMP plays a key role in controlling cytoplasmic turgor and cell volume through mechanosensitive osmolyte channels (67). Cells respond to c-di-AMP buildup by reducing osmolyte uptake and activating K^+^ export (68), thus decreasing influx of water and cellular volume (69). Our observation that EpaU binds generally over the *E. faecalis* cell surface but does not alter cell morphology or stimulate autolysis in the absence of a membrane-permeabilizing agent suggests that EpaU may function as a remodeling peptidoglycan hydrolase by making the peptidoglycan meshwork more porous and less stiff, enabling cell wall enlargement. We propose that EpaU-induced changes in the peptidoglycan meshwork are sensed by an unknown mechanism that adjusts c-di-AMP levels, activating osmolyte uptake and water influx. Interestingly, deletion of AtlA, a peptidoglycan hydrolase which plays a predominant role in cell separation, or treatment of bacteria with lysozyme, does not alter c-di-AMP levels in *E. faecalis*, implying that this sensing mechanism recognizes specific signals generated by EpaU cleavage. An increase in internal osmotic pressure, promoted by EpaU activity, would lead to cell expansion, pushing the plasma membrane against the peptidoglycan layer. There is now ample experimental evidence in different bacteria that c-di-AMP positively modulates cell wall thickness, and β-lactam resistance and tolerance (8–11). For example, in *Lactococcus lactis*, accumulation of c-di-AMP correlated with elevated levels of peptidoglycan precursor UDP-GlcNAc (70), suggesting a direct connection between c-di-AMP synthesis and peptidoglycan biosynthesis. However, we observed that increased c-di-AMP levels in the WT::p*cdaA* strain do not reduce sensitivity to ampicillin. Thus, the reason for the reduced sensitivity of Δ*epaU* to ampicillin is currently not clear, but it could be due to the participation of EpaU in the cell lysis induced by ampicillin. Further exploration of EpaU’s role in ampicillin resistance is required to validate this hypothesis.

We further showed that EpaU binding to the TA decorations of EPA is crucial for efficient lysis of *E. faecalis* cells. How EpaU activity is regulated to prevent adventitious lysis of the bacteria under normal growth conditions is unclear. Of note, the N-terminal domain of *E. faecalis* autolysin AtlA was previously reported to be glycosylated, and its processing was required for AtlA-mediated septum cleavage (71). Intriguingly, similarly to AtlA, the EpaU N-terminal domain is a serine/threonine-rich intrinsically disordered region. We recently reported that such regions in extracellular proteins of three streptococcal species are glycosylated (51). The cell surface-associated EpaU is predominantly secreted as two forms, suggesting O-glycosylation of EpaU at the N-terminal domain. Future studies will examine whether this domain controls the EpaU activity.

In summary, our study demonstrates the crucial role of TA decorations in binding the CWB domain of EpaU and provides novel insights into the V583 EPA structure, as well as the roles of EpaR and EpaX in the synthesis of TA decorations. Further work is needed to elucidate the structure of EPA in V583 and other strains. This analysis will be critical for investigating how EPA is synthesized and surface presented, and how it interacts with other *E. faecalis* autolysins and peptidoglycan remodeling enzymes. Our data also point to a previously unrecognized link between c-di-AMP signaling and peptidoglycan cleavage in *E. faecalis*. Understanding this mechanism could help to identify sensors that monitor the integrity and mechanical properties of the peptidoglycan meshwork, offering novel strategies to address the growing threat of *E. faecalis* infections.

## Methods

### Bacterial strains, growth conditions and media

All strains, plasmids, and primers used in this study are listed in Tables S3 and S4. *E. faecalis* strains were grown in BD Bacto Todd-Hewitt broth supplemented with 1% yeast extract (THY) without aeration at 37 °C with 5% CO_2_. Unless otherwise indicated, frozen glycerol stocks of *E. faecalis* were prepared from overnight-grown strains resuspended in THY+20% glycerol. *E. coli* strains were grown in Lysogeny Broth (LB) medium or on LB agar plates at 37 °C. When required, antibiotics were included at the following concentrations: chloramphenicol at 10 µg mL^-1^ for *E. coli* and *E. faecalis*; erythromycin at 300 µg mL^-1^ for *E. coli* and at 5 µg mL^-1^ for *E. faecalis*; kanamycin at 50 µg mL^-1^ for *E. coli*.

### Construction of *E. faecalis* strains

*E. faecalis* mutant strains were constructed using homologous recombination using the pLT06 vector as described (53). Plasmids for mutagenesis were constructed by PCR amplifying ∼1,000 bp fragments upstream and downstream of the gene of interest and ligating them into the pLT06 vector. To construct complemented strains for Δ*epaR* and Δ*epaX* strains, the DNA fragments corresponding to *epaR* and *epaX* genes were PCR-amplified and cloned into the pDC123 vector for constitutive expression. To construct a complemented strain for Δ*epaU*, the *epaU* gene was amplified and ligated into the pMSP3535 vector for nisin-inducible expression (50). Plasmids for expression of the catalytic mutant EpaU E262Q (p*epaU* E262Q) and a truncated EpaU lacking the cell-wall binding domain (p*epaU*ΔCWB) were generated using site-directed mutagenesis. pMSP3535 was a gift from Gary Dunny (Addgene plasmid # 46886). Plasmids for expression of *cdaA* encoding the diadenylate cyclase DacA and *yycJ* encoding a protein of unknown function (72) (negative control) were constructed using the pMSP3535 vector. All plasmids were verified by Sanger sequencing. The complementation and control plasmids were introduced into the WT and mutant strains by electroporation.

### Construction of the *E. coli* expression plasmids for recombinant production of EpaU, GFP-EpaU^V583^ and GFP-EpaU^OG1RF^

To construct vectors for expression of full-length EpaU and the C-terminal fragment of EpaU, the DNA fragments were PCR-amplified and ligated into the pRSF-NT vector that contains the N-terminal His_6_-tag followed by a TEV protease cleavage site. To construct a vector for expression of GFP-EpaU^V583^ fusion, the DNA fragments corresponding to superfolder GFP and the C-terminal domain of EpaU^V583^ were PCR amplified, assembled using NEBuilder HiFi master mix (NEB), and ligated into the pRSF-NT vector. The vector for expression of GFP-EpaU^OG1RF^ fusion was constructed similarly, except that the high sequence similarity between CWB domains of EpaU^OG1RF^ precluded PCR amplification from *E. faecalis* OG1RF genomic DNA. An optimized synthetic DNA fragment corresponding to the C-terminal domain of EpaU^OG1RF^ (Blue Heron) was used for PCR amplification and subsequent cloning into the pRSF-NT vector.

### Protein expression and purification of EpaU, GFP-EpaU^V583^ and GFP-EpaU^OG1RF^

The EpaU, GFP-EpaU^V583,^ and GFP-EpaU^OG1RF^ construct plasmids were transformed into *E. coli* LOBSTR BL21(DE3) cells, and a starter culture containing 2% glucose in LB was grown overnight. A 1:60 dilution of the overnight culture was inoculated into LB, and the culture was grown to an OD_600_ of 0.6 at 37 °C before the addition of 400 µM Isopropyl β_-_D_-_thiogalactopyranoside (IPTG) for induction at 16 °C. After overnight induction for 12 h, cultures were centrifuged at 4,302 g for 20 min at 4 °C. Bacteria were resuspended in a buffer of 20 mM Tris pH 8.0, 300 mM NaCl, 10 mM imidazole.

Bacteria were lysed using two passes through a Microfluidizer (Microfluidics). Cell debris was removed via centrifugation at 38,758 g for 1 h at 4 °C. EpaU variants were purified by passage over a Ni-affinity column containing His-Trap chelating resin from GE Healthcare Life Sciences. The column was washed with a buffer of 20 mM Tris pH 8.0, 300 mM NaCl, 10 mM imidazole. EpaU variants were then eluted using the same buffer with 250 mM imidazole. After assaying eluates for purity by SDS-PAGE, fractions containing the protein of interest were spin-concentrated using a Amicon Ultra 4 with a 3,000 MWCO (Millipore) and then buffer exchanged into a final storage buffer of 20 mM Tris pH 8.5, 100 mM NaCl using continuous dilutions in the spin concentrator and flash frozen in liquid nitrogen.

### Analysis of EpaU expression in *E. faecalis*

Starting from frozen bacterial stocks, 40 mL of *E. faecalis* culture was grown in THY to an OD_600_ of 0.55. Cells were then centrifuged at 3,124 g, washed with 30 mL of phosphate-buffered saline (PBS), and resuspended in 250 µL of PBS containing 2% SDS. Samples were then gently rotated at room temperature for 1 h before centrifugation at 3,200 g for 10 min. Immunoblotting was then performed on a PVDF membrane with rabbit polyclonal antibodies recognizing the C-terminal binding domain of EpaU (1:500 dilution). Rabbit polyclonal antibodies against the C-terminal fragment of EpaU were produced by Cocalico Biologicals. Horseradish Peroxidase-conjugated secondary goat anti-rabbit antibodies (ThermoFisher Scientific, 32460) at a dilution of 1:2,500 were used for signal detection on a ChemiDoc imaging system (BioRad).

### Analysis of proteins in a cell-free culture supernatant of *E. faecalis*

Cultures were grown from frozen stocks to an OD_600_ of 0.55 and then centrifuged at 16,000 g for 2 min. As a positive control for cytosolic content leakage, WT cells were lysed with disruptor beads (EMS, d=0.1mm) at 4 °C for 15 min or incubated with 0.5 mg mL^-1^ lysozyme. After filtration through a d=0.2 µm syringe filter, supernatants were used for TCA protein precipitation as described with modifications (73). Briefly, cold 20% TCA was added to the supernatants, and the mixture was incubated on ice for 30 min. Supernatants were then centrifuged at 18,213 g for 15 min at 4 °C. Pellets were washed twice with cold 100% acetone, dehydrated, and resuspended in SDS sample buffer. Proteins were separated on 10% Bis-Tris gel, transferred onto a nitrocellulose membrane, and probed with primary anti-EpaU (1:1,000) and secondary goat-anti-rabbit-HRP (1:10,000) antibodies. After 30 min incubation with stripping solution (1.5% Glycine, 1% SDS, 1% Tween-20 at pH 2.2), the membrane was re-probed with anti-HSP60 mAb (Abcam, 1:1,000) and goat-anti-mouse-HRP (1:10,000) before detection in a ChemiDoc imaging system (BioRad).

### Fluorescent and differential interference contrast (DIC) microscopy

The exponential phase bacteria (OD_600_ of 0.3) were fixed with paraformaldehyde (4% final concentration) and pipetted onto microscope slide cover glasses (high performance, D=0.17 mm, Zeiss) coated with poly-L-lysine, and allowed to settle for one hour at room temperature. Bacteria were incubated with GFP-EpaU^V583^at a final concentration of 1.56 µg mL^-1^, 15.6 µg mL^-1^or 156 µg mL^-1^for 15 min at room temperature. The samples were washed four times with PBS and mounted on a microscope slide with ProLong Glass Antifade Mountant (Invitrogen). Samples were imaged on a Leica SP8 equipped with a 100×, 1.44 N.A. objective and DIC optics. Images were deconvolved using Huygens Professional and ImageJ software.

### Scanning electron microscopy

The exponential phase bacteria (OD_600_ of 0.7) were fixed with paraformaldehyde (4% final concentration) and then pipetted onto microscope slide cover glasses (12 mm Circle No. 2) coated with poly-L-lysine. Following 1 h incubation at room temperature, the cover glasses were washed 3 times with PBS. Bacteria were dehydrated stepwise in a gradient series of ethanol (35%, 50%, 70%, 80%, and 96% for 20 min each, and then in 100% overnight at −20 °C), followed by critical point drying with liquid CO_2_ in a Leica EM CPD300. Samples were coated with 5 nm of platinum. SEM images were performed in immersion mode on an FEI Helios Nanolab 660 dual-beam system using a TLD detector and secondary electron mode.

### Autolysis assay

WT (*E. faecalis* strain VE14089) and the Δ*epaU* mutant were grown overnight in THY at 37 °C. The next day, strains were inoculated at a 1:20 dilution into 40 mL of fresh THY and grown at 37 °C until OD_600_ reached 0.5. Cultures were then centrifuged at 3,900 g for 10 min and washed with 10 mL of PBS. Cultures were then centrifuged again and resuspended in 10 mL of fresh PBS. 100 µL of bacterial suspension was added to a well of a 96-well plate with 100 µL of PBS containing various concentrations of Triton X-100. Autolysis was measured in a Cytation 7 (Biotek) plate reader at 600 nm for 4 h at 37 °C, with measurements taken every 15 min with the plate shaken before each measurement.

### Bactericidal assay with exogenous EpaU

To prepare frozen stocks, WT (VE14089) and Δ*epaR* were grown to mid-exponential phase (OD_600_ of 0.5), collected by centrifugation at 3,900 g for 10 min, washed with the buffer (20 mM HEPES pH 7.5, 2 mM CaCl_2_, 1% BSA), and then frozen in the buffer containing 15% glycerol. Stocks were diluted to 800 CFU for the experiment. 200 µL of the bacterial culture was added to a 96-well plate along with 10 µL of EpaU at varying concentrations, and the mixture was incubated for 2 h at 37 °C. After incubation, 50 µL of bacterial suspension from each well was plated onto THY agar plates and incubated overnight at 37 °C. CFUs were enumerated the next day and expressed as a percentage of survival versus the control well containing no EpaU.

### Saccule binding

Bacterial saccules were prepared by growing 45 mL of *E. faecalis* strains in THY to an OD_600_ of 0.55. After centrifugation at 3,124 g and resuspension in PBS, cells were spun again, resuspended in 700 μL of PBS, spun at 23,447 g, resuspended in 1 mL of 4% SDS, and incubated for 1 h at 100 °C. Saccules were then washed with 1 mL of 1M NaCl four times after centrifugation at 23,447 g, followed by four additional washes with distilled water. Saccules were resuspended in PBS to an OD_600_ of 3.0. 500 μL of saccules was combined with 0.05 mg of GFP-tagged protein and incubated for 1 h at room temperature with rotation. 100 μL of the sample was added to a black-bottom microplate, and the total fluorescence of the sample was read using a SpectraMax M5 (Molecular Devices) with an excitation at 485 nm and emission at 510 nm. The sample was returned to the initial volume. Saccules were then spun down and washed three times with 1 mL of PBS, then resuspended to a final volume of 500 μL plus the volume of the protein delivered. Another 100 μL aliquot of the sample was removed, and fluorescence was measured again to obtain the bound signal. The percentage of fluorescence remaining was then calculated from the residual signal.

### EPA release and purification Isolation of cell walls

Cell walls were isolated from the exponential phase cultures (OD_600_ of 0.7) by the SDS-boiling procedure as described for *S. pneumoniae* (74). Purified cell wall samples were lyophilized and stored at −20 °C before the analysis.

### Purification of mild acid released EPA for chromatographic analysis (Fig. 5A)

EPA was released from the purified cell wall of *E. faecalis* strains by mild acid hydrolysis (0.02 N HCl at 100 °C for 20 min) as previously described for *S. pyogenes* cell wall polysaccharide (75). Supernatant containing the released EPA was then concentrated and desalted by spin column (Amicon 3,000 MWCO filters) to a volume of 200 µL and fractionated on a column of BioGel P150 (18 mL, equilibrated and eluted in 0.2 N sodium acetate, pH 3.7, 0.15 M NaCl, collecting fractions of 0.7 mL). Samples containing polysaccharides were determined by an anthrone assay as described below. These fractions were again combined, concentrated, and desalted using a spin column (Amicon 3,000 MWCO filters) and resolved by anion-exchange chromatography on an 18 ml column of DEAE Toyo 650M equilibrated in 5 mM HEPES-NaOH, pH 7.4. The columns were eluted with 40 mL of starting buffer and then with a gradient (80 mL) of NaCl (0-0.2 M, dashed line). Fractions were analyzed for carbohydrate by anthrone assay, phosphate by malachite green assay, and subjected to ANDS tagging and electrophoresis as described below.

### Purification of enzymatically released EPA for chromatographic analysis (Fig. 6A)

To release EPA from peptidoglycan by enzymatic hydrolysis, 5 mg mL^-1^ of the purified cell wall material was resuspended in the lysis buffer containing 5 U mL mutanolysin (Sigma-Aldrich), 50 µg mL^-1^ zoocin A, 20 mM Tris-HCl pH 7.5 and 300 mM NaCl. Following overnight incubation at 37 °C, soluble polysaccharides were separated from the cell wall by centrifugation at 13,000 g, 10 min. The supernatant was concentrated and desalted using a spin column (Amicon 3,000 MWCO filters) to a volume of 200 µL and fractionated on a column of BioGel P150 (18 mL, equilibrated and eluted in 0.2 N sodium acetate, pH 3.7, 0.15 M NaCl, collecting fractions of 0.7 mL). Fractions containing EPA, as determined by an anthrone assay, were combined and desalted on a PD-10 desalting column (Cytiva). This sample was then re-concentrated using a spin column (Amicon 3,000 MWCO filters) and hydrolyzed by mild acid (0.02 N HCl at 100 °C for 20 min). Mild acid-hydrolyzed EPA was then concentrated and desalted by spin column (Amicon 3,000 MWCO filters) and resolved by anion exchange chromatography on a 5 mL HiTrap DEAE FF Fast Flow column (Cytiva). Fractions were assayed for carbohydrate by anthrone assay, phosphate by malachite green assay, and subjected to ANDS tagging and electrophoresis as described below.

### Purification of enzymatically released EPA for GC-MS (Tables 1 and 2)

For GC-MS experiments, the soluble fraction of enzymatically released EPA was precipitated with 80% acetone at −20 °C overnight. Precipitated EPA was recovered by centrifugation at 3,200 g, 20 min. The acetone pellets were dried briefly under a stream of air, dissolved in 0.2 mL of water, and fractionated on a column of BioGel P150 (18 mL, equilibrated and eluted with 0.2 N sodium acetate, pH 3.7, 0.15 M NaCl, collecting 0.7 mL fractions).

### ANDS labeling of EPA fractions

Fractions of purified EPA from DEAE columns were fluorescently labelled with ANDS by reductive amination as previously described (21, 76). After amination with ANDS, samples were desalted by repetitive cycles of dilution and concentration on a 3,000 MWCO spin filter (Amicon) five times with 0.4 mL of water and recovered in a final volume of 0.2 mL. Fluorescent EPA fractions were analyzed by SDS-PAGE on 8-12% SurePage gradient gels as described previously (21, 76) and imaged on a BioRad ChemiDoc imager.

### Anthrone assay

Rhamnose and hexose contents were measured by anthrone assay as previously described (21, 76). Briefly, the hexose content was measured directly from the absorbance at 700 nm, employing a glucose standard curve for comparison. The rhamnose content was determined at 580 nm, following subtraction of the contribution from hexose, determined from the absorbance at 700 nm. The absorbance at 580 nm of the hexose chromophore is approximately equivalent to the absorbance at 700 nm, whereas the contribution of rhamnose to absorbance at 700 nm is negligible. To monitor rhamnose recoveries during routine isolation and purification procedures, rhamnose content was estimated by anthrone analysis at 580 nm. A standard curve of L-rhamnose was prepared for quantification.

### Phosphate assay

Phosphate content of EPA fractions was measured by the malachite green method as previously described (75). Briefly, 20 µL of each fraction was treated with 2M HCL for 1 h at 100 °C, then neutralized with 1 eq. of NaOH. Samples were cooled to room temperature and brought to a final volume of 100 µL with buffer (HEPES, pH 7.5). 100 units of alkaline phosphatase (New England Biolabs) were added along with 10 µL of the provided alkaline phosphatase buffer. Samples were rotated overnight at 37 °C. The next day, 100 µL of the sample was added to a 96-well plate with 200 µL of malachite green reagent (1 vol 4.2% ammonium molybdate tetrahydrate (by weight) in 4 M HCl, 3 vol 0.045% malachite green (by weight) in water, and 0.01% Tween 20). The plate was incubated for 10 min, and the absorbance was measured at 620 nm. A standard curve for phosphate was generated alongside the samples to determine their concentrations.

### Analysis of glycosyl composition by gas chromatography-mass spectrometry (GC-MS)

Composition of partially purified EPA was determined as trimethylsilyl (TMS) derivatives of O-methyl glycosides following methanolysis in methanol/1 N HCl (100 °C, 3 h) either in-house as described (75) or at the Complex Carbohydrate Research Center (CCRC), University of Georgia, Athens, GA (77). In some analyses, samples were preincubated overnight with HF to improve ribitol detection by removing phosphate. Samples were dissolved in 0.025 mL 51% HF, incubated at 4 °C overnight, dried thoroughly under a stream of nitrogen gas, dissolved in water, transferred to a 13×100 screw cap tube, dried again under nitrogen, and treated as above for GC-MS analysis.

Reducing-end analysis was performed on the mild acid-released EPA material as alditol acetates (78) following sodium borohydride reduction (10 mg mL^-1^NaBH_4_, 0.1 M NH_4_OH). After overnight chemical reduction at room temperature, reactions were neutralized with acetic acid and chromatographed on BioGel P150, as described above. Reduced EPA was recovered, desalted, and hydrolyzed in 2 N trifluoroacetic acid at 120 °C for 2 h. Samples were evaporated out of isopropanol under a stream of air three times, per-acetylated in 0.2 mL pyridine/acetic anhydride (1:1) at 100 °C for 1 h, and diluted with 3 mL of CHCl_3_. Reactions were partitioned 2 times with 2 mL of water, dried gently under a stream of air, and analyzed by GC-MS on a Thermo Scientific Trace 1310 gas chromatograph, equipped with a 0.25 mm x 15 m DB-1701 (J&W Scientific) capillary column, as previously described (75).

### Analytical ultracentrifugation (AUC)

Sedimentation velocity analytical ultracentrifugation experiments were performed using a Beckman Coulter ProteomeLab XL-I. Absorbance optics were utilized at 280 nm. An 8-hole rotor was used to spin all four samples in the same experiment. Samples were prepared by incubating 1 mg mL^-1^ of GFP-EpaU^V583^ protein with EPA polysaccharide samples for 20 min on ice. The concentration of EPA polysaccharide was calculated based on the rhamnose concentration. The final concentrations of rhamnose were 4.55 mM for WT, 7.51 mM for Δ*epaX*, and 8.58 mM for Δ*epaR* for enzymatically released EPA samples; 4.22 mM for WT, 4.91 mM for Δ*epaX*, and 6.96 mM for Δ*epaR* for mild-acid released EPA samples. Standard Beckman Coulter 2-channel Epon charcoal centerpieces were loaded with approximately 450 µL of sample and buffer. The buffer used was 20 mM HEPES, 150 mM NaCl, pH 7.5. Data were collected at 20 °C until protein sedimentation was complete. The wide distribution analysis (WDA) was performed using all scans collected and was conducted in SEDANAL (v7.86) (79, 80). SEDFIT (v16.50) was used for the continuous c(*s*) distribution analysis (81, 82). The buffer density and viscosity were estimated using SEDNTERP (v3.0.4) (83). The partial specific volume was set to 0.73 mL/g. Normalization by area of the WDA distributions was performed directly in SEDANAL for the region from 1 – 20 S. The normalization of c(*s*) distributions by area was performed in Excel using the integrated areas between 1-15 S reported by GUSSI (v2.1.2) (84).

The standardization of the sedimentation coefficient from apparent (*s*\*) to that of water at 20 °C (S_20,w_) was calculated according to the following equation,

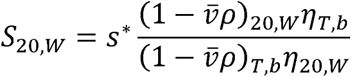

where v is the partial specific volume (mL/g), p is the buffer density (g/mL), and) is the buffer viscosity (centipoise; cp). The subscript, b specifies that the parameter is for the experimental buffer and temperature, whereas the subscript 20, W specifies the standard condition at 20 °C in water (80).

### Antibiotic resistance assays

The bacterial growth was initiated by a 1:100 dilution of starting cultures with an OD_600_ of 0.55 in THY, followed by incubation of microdilution plates at 37 °C for 12 h, after which the final optical density was measured using a BioTek Epoch 2 microplate spectrophotometer. MIC values were determined in triplicate for each strain using a modified Gompertz function as previously described by Lambert and Pearson to calculate MICs and confidence intervals (CI) directly (85).

### Quantification of intracellular c-di-AMP

Quantification of intracellular c-di-AMP was performed using the Cyclic di-AMP ELISA kit from Cayman Chemicals (Item No. 501960). Samples were prepared by growing 10 mL of bacterial cultures (OD_600_ of 0.55) and then spinning down at 3,124 g, followed by washing with 30 mL of PBS. The cell pellet was resuspended in 800 µL of ice-cold assay buffer (100 µM Tris pH 7.5, 20 mM MgCl_2_), 50 mM NaCl with 10 µL of 25 kU mutanolysin. Cells were lysed using 50 µL of 0.1 mm glass disruptor beads (Electron Microscopy Sciences) via bead-beating for 1 h at room temperature in a vortex shaker. Beads were then allowed to settle for 20 min as the cells were cooled to 4 °C, after which the lysate was decanted. Lysate was then heated to 95 °C for 10 min to stop all enzymatic activity. Lysate was spun again to remove precipitated proteins and any remaining glass disruptor beads. A pilot ELISA suggested that a 2:1 dilution placed the c-di-AMP concentration within the linear range of the assay for all samples except Δ*epaU* and its empty vector control, Δ*epaU*::pMSP3535, which were diluted at a 4:1 ratio using assay buffer. Samples were then used as unknowns in the Cayman ELISA kit following all the suggested protocol, with each plate experiment using its own calibration curve. The plate was read at 450 nm on a SpectraMax M5 (Molecular Devices) after development. Blank subtraction, calibration curve fitting, and interpolation of unknowns were performed using GraphPad Prism 10 software. Absorbance of all standard curves was within ±5%. For the peptidoglycan challenge assay, lysozyme was added to cultures at the start of growth. Prior to the experiment, growth analysis was performed using 0.25-20 mg mL^-1^ of lysozyme to determine the concentrations affecting *E. faecalis* fitness. Concentrations 0.25-0.75 mg mL^-1^ of lysozyme did not affect *E. faecalis* growth and were used to measure c-di-AMP concentrations.

### Statistical analysis

Unless otherwise indicated, statistical analysis was carried out on pooled data from at least three independent biological repeats. Statistical analyses were performed using Graph Pad Prism version 10.2.0. Quantitative data was analyzed using one-way ANOVA, 2-way ANOVA, and unpaired t-test or Dunnett’s multiple comparisons test as described for individual experiments. A *P*-value equal to or less than 0.05 was considered statistically significant.

## Acknowledgments

The authors thank Dr. Catalina Velez-Ortega, University of Kentucky, for access to the Leica SP8 confocal microscope. We thank Lynn Hancock, University of Kansas, for providing the pLT06 plasmid and the Δ*atlA* and Δ*cpsC* strains; Julia Willett, University of Minnesota, for providing the OG1RF strain.

Scanning electron microscopy was performed at the Electron Microscopy Center (University of Kentucky), member of the KY INBRE (Kentucky IDeA Networks of Biomedical Research Excellence), which is funded by the National Institutes of Health (NIH) National Institute of General Medical Sciences (IDeA Grant P20GM103436), which belongs to the National Science Foundation NNCI Kentucky Multiscale Manufacturing and Nano Integration Node, supported by ECCS-1542174. Carbohydrate composition/linkage analysis at the Complex Carbohydrate Research Center was supported by the U.S. Department of Energy, Office of Science, Basic Energy Sciences, Chemical Sciences, Geosciences and Biosciences Division, under award DE-SC0015662 NIH grant R24GM137782 to P.A. This work was supported by NIH grant R21AI166233 from the NIAID to K.V.K. The content is solely the responsibility of the authors and does not necessarily represent the official views of the National Institutes of Health.

## Author contributions

J.S.R., C.T.C., N.K., and K.V.K. designed the experiments. N.R.M., J.S.R., L.H., C.T.C., and C.W.K. performed functional and biochemical experiments. S.Z. performed microscopy analysis. A.E.Y., A.B.H., and C.T.C. performed and analyzed analytical ultracentrifugation experiments. K.V.K. constructed plasmids and isolated mutants. N.R.M., C.T.C., J.S.R., S.Z., P.A., K.V.K., and N.K. analyzed the data. N.K. and C.T.C. wrote the manuscript with contributions from all authors. All authors reviewed the results and approved the final version of the manuscript.

## Data availability

All data generated during this study are included in the article and Supplementary Information files.

