## Supplemental Material for "*Enterococcus faecalis* autolysin, EpaU, binds the enterococcal polysaccharide antigen via its teichoic acid-like repeats and modulates c-di-AMP signaling"

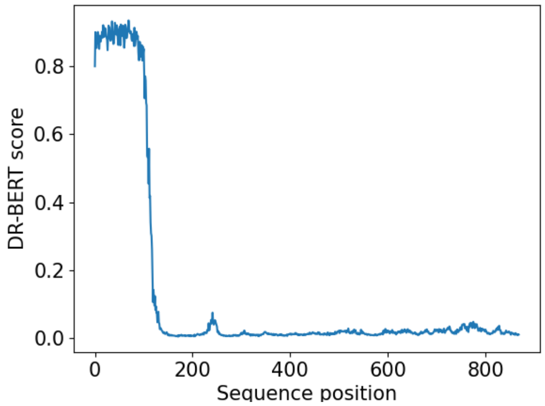


Figure S1. **The protein disorder prediction of EpaU$^{V583}$ using DR-BERT.**
The protein disorder was predicted using a protein language model DR-BERT (1). Values above 0.75 are indicative of protein disorder. The signal peptide sequence is omitted.

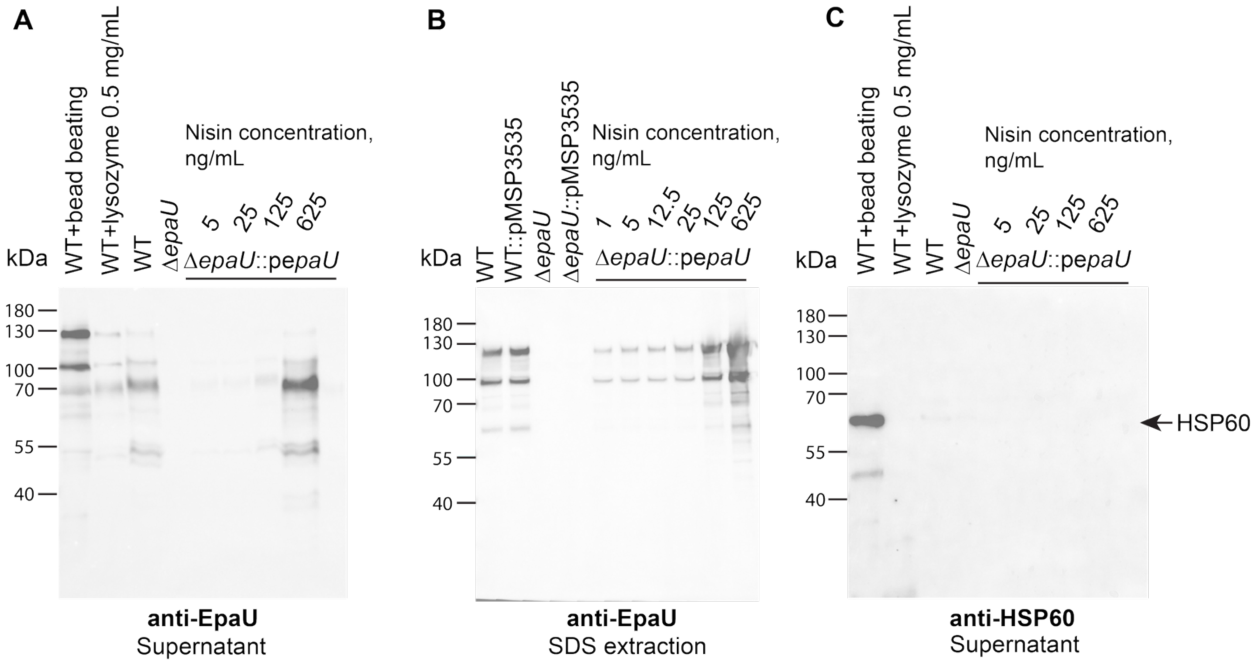


Figure S2. **Analysis of EpaU expression and the presence of HSP60 in the cell-free supernatant.**

(**A, B**) Immunoblots of EpaU precipitated from culture supernatants (A) and extracted from the cell surface with 2% SDS (B). (**C**) Proteins precipitated from culture supernatants were probed for heat-shock protein 60 (HSP60) using anti-HSP60 antibodies. EpaU expression in Δ*epaU*::p*epaU* was induced with varying concentrations of nisin. WT cells lysed by bead-beating or treated with lysozyme (0.5 mg $mL^{-1}$) were used as positive controls for HSP60 analysis. The experiments were performed independently two times and yielded the same results. A representative image from one experiment is shown.

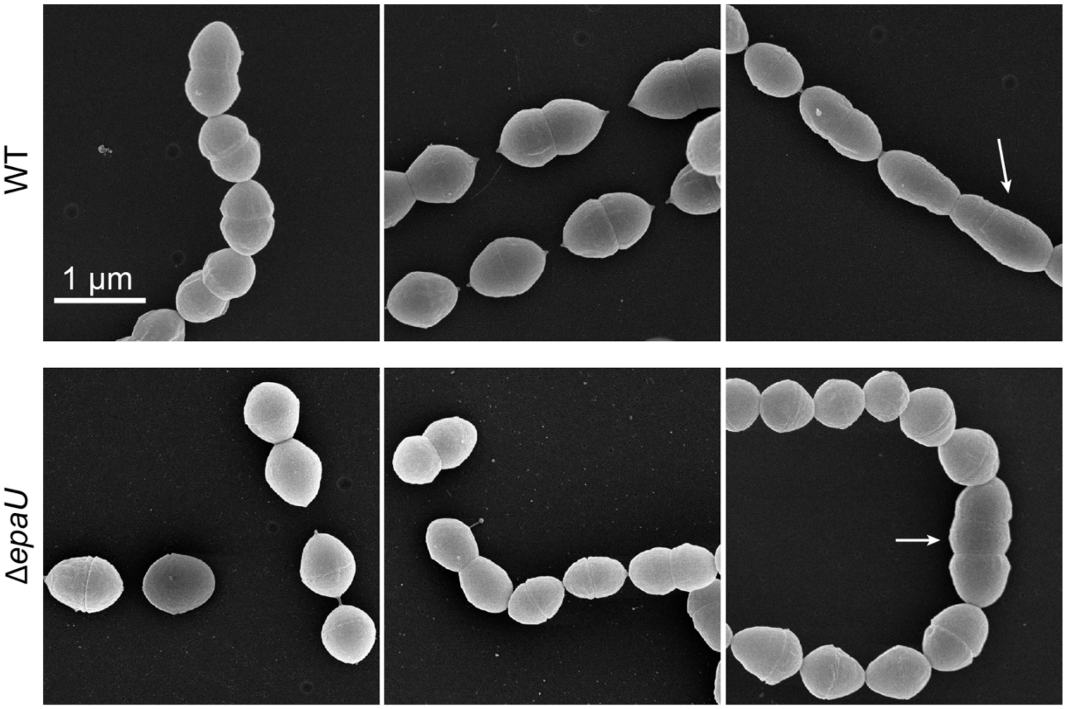


Figure S3. **Scanning electron micrographs of *E. faecalis* strains.**
Exponential phase bacterial cells were fixed, dehydrated, coated with platinum, and visualized by scanning electron microscopy. The majority of the cells in both strains are oval-shaped and connected into chains. White arrows denote cells with irregular cell division present in the WT and Δ*epaU* cells with similar frequency: 4.73% for WT (n=148) and 5.06% for Δ*epaU* (n=158), where n = number of cells observed. The imaging was performed independently three times and yielded the same results. Scale bar – 1 µm.

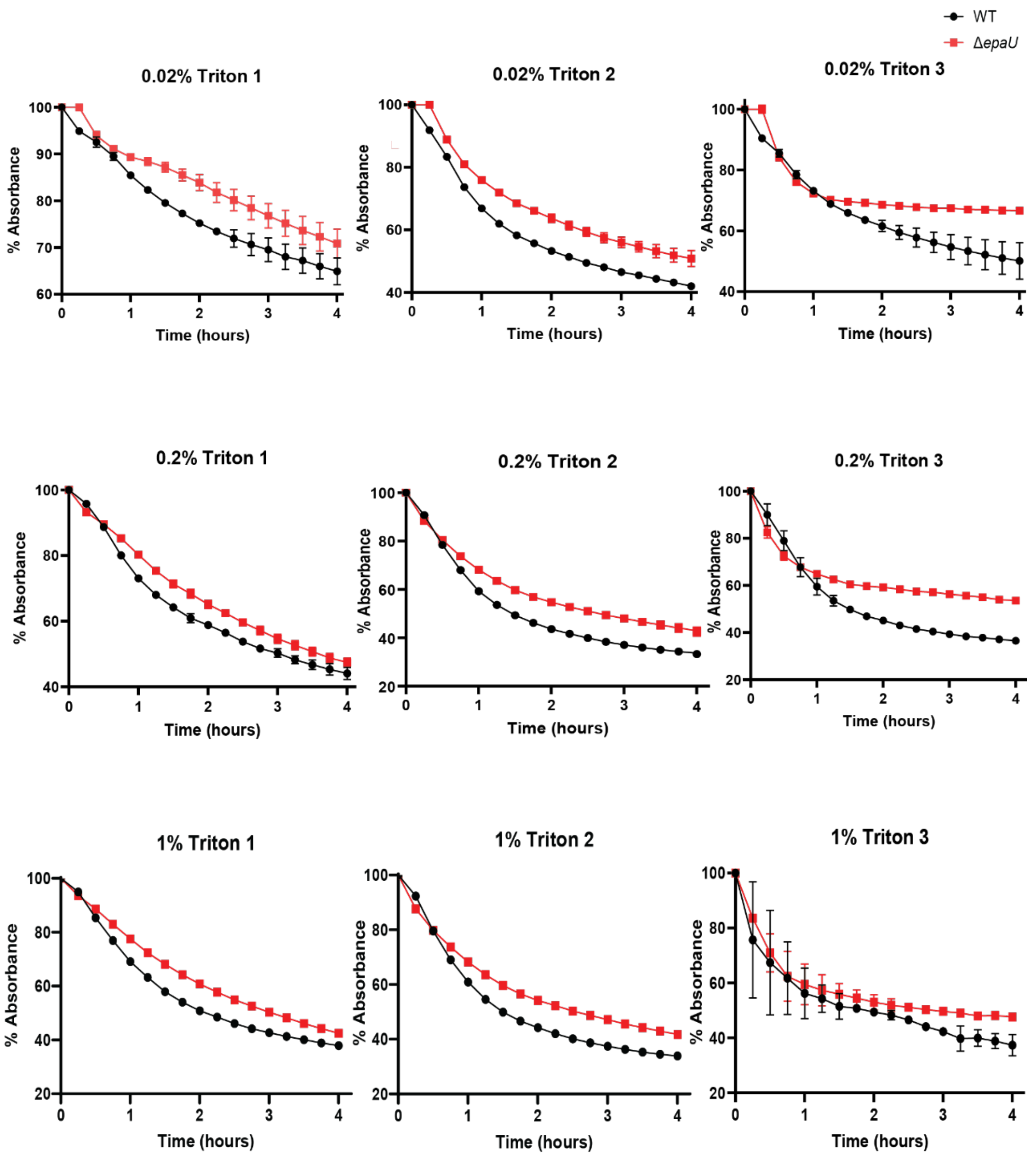


Figure S4. **Autolysis of WT (black line) vs Δ*epaU* (red line).**
Each biological replicate is shown at all tested Triton X-100 concentrations. Autolysis was monitored as the decrease in $OD_{600}$ and normalized to the $OD_{600}$ at time zero (100%). The error bars represent standard errors between technical repeats.

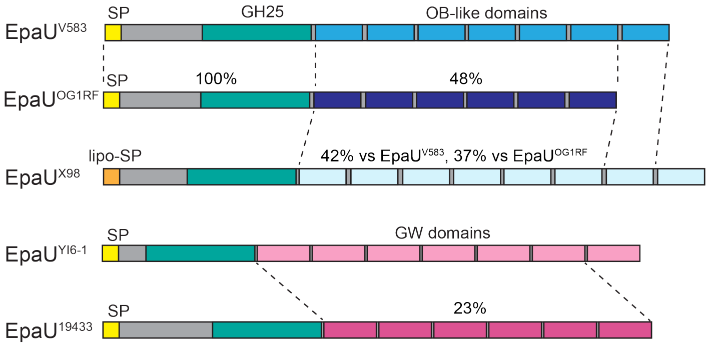


Figure S5. **The domain diagrams of EpaU autolysins from different *E. faecalis* strains.**

While all EpaU autolysins possess endo-N-acetylmuramidases catalytic domains (GH25 family), their cell wall-binding domains differ. The cell wall-binding domains of EpaU from V583, OG1RF and X98 strains have a variable number of repeat OB-like (Oligonucleotide/Oligosaccharide-Binding) domains (DUF5776) with limited sequence identity indicated by percentages. The cell wall-binding domains of EpaU from YI6-1 and ATCC 19433 strains have a variable number of repeat GW (Gly-Trp dipeptide) domains (PFAM 13457) with low sequence identity to each other and no homology to the structurally unrelated OB-like domains of EpaU from V583, OG1RF and X98 strains.

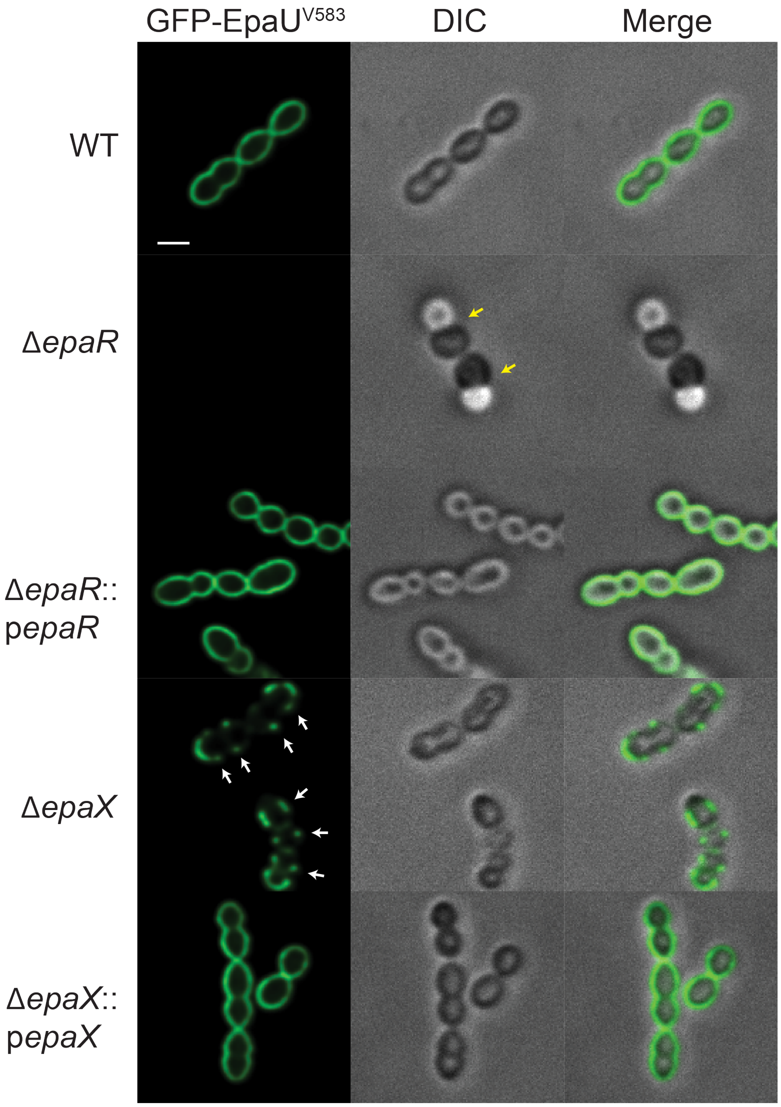


Figure S6. **Binding of 15.6 μg mL$^{-1}$ GFP-EpaU$^{V583}$ to *E. faecalis* cells.**
Exponential phase cells of the WT, Δ*epaR*, Δ*epaR*::p*epaR*, Δ*epaX* and Δ*epaX*::p*epaX* strains were incubated with 15.6 μg mL$^{-1}$ GFP-EpaU$^{V583}$ and examined with differential interference contrast (DIC) and fluorescent microscopy. White arrows indicate equatorial sites labeled by GFP-EpaU$^{V583}$. Yellow arrows indicate cell division defects. The experiments were performed independently three times and yielded the same results. A representative image from one experiment is shown. Scale bar is 1 μm.

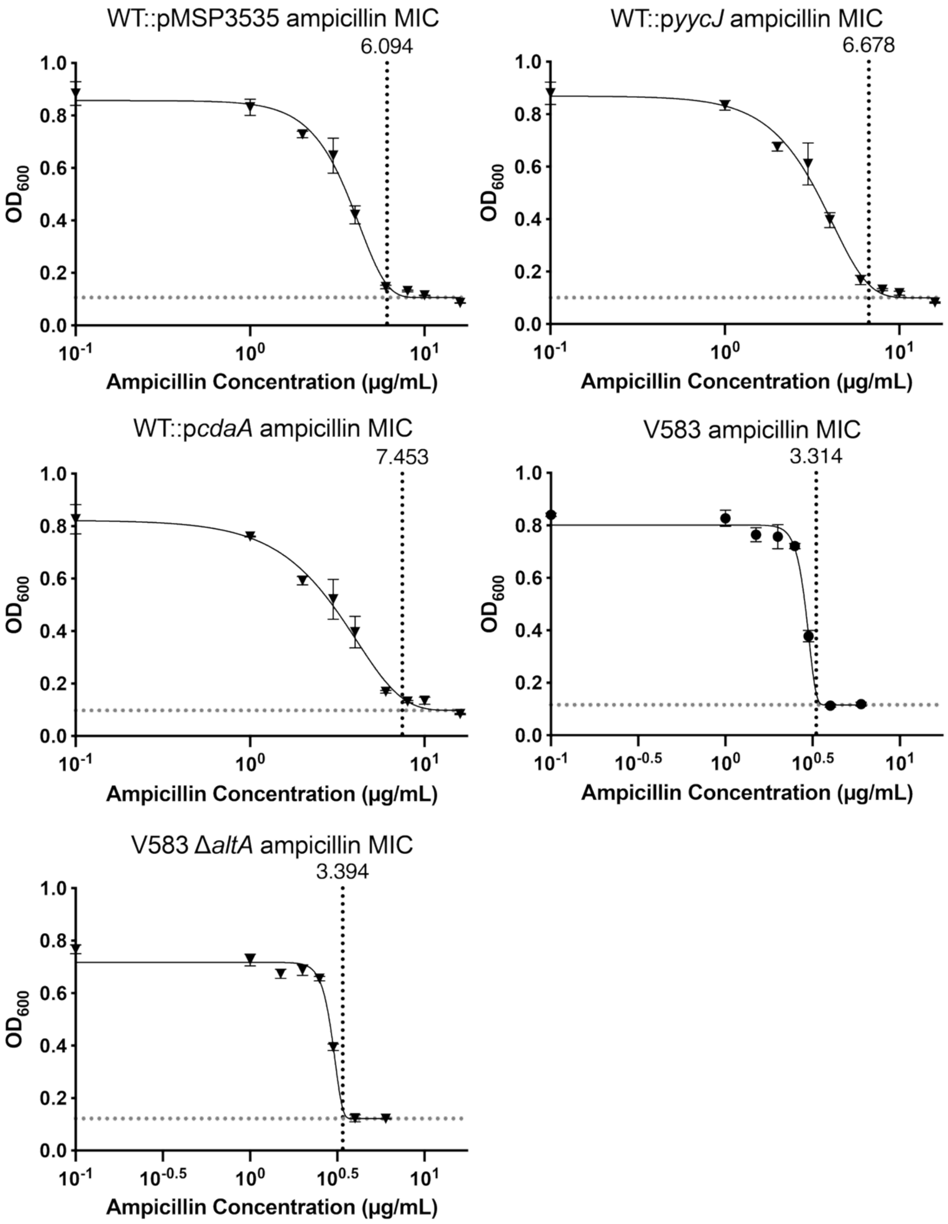


Figure S7. Best-fit curves for ampicillin resistance controls of wild-type expressing the empty pMSP3535 vector, and wild-type over-expressing both *yycJ* and *cdaA*. The WT::p*yycJ* was used as an additional control. The experiments with WT::pMSP3535, WT::p*yycJ*, and WT::p*cdaA* strains were performed with 25 ng/ml nisin added as an inducer. The Δ*altA* mutant was also measured alongside its V583 parent strain for reference. The vertical dotted line represents the best-fit MIC. MIC values are reported in μg/mL and were computed using Gompertz equation for MIC determination (2). Fits are the result of three independent biological repeats.

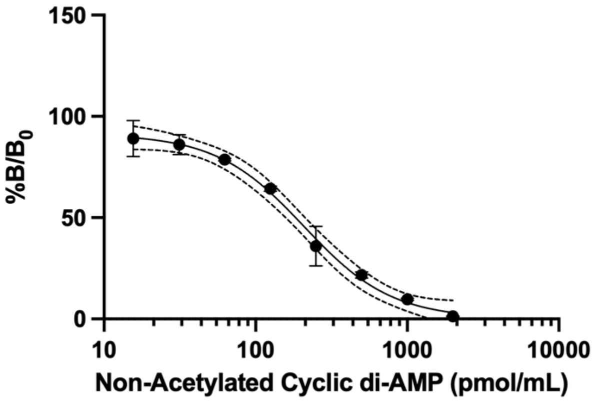


Figure S8. **Example standard curve for non-acetylated cAMP**.
Interpolation of unknowns was performed using an experimental standard curve measuring bound/maximum bound (%B/$B_0$) to known cAMP concentration for each assay run. 95% confidence intervals are shown as dashed lines using a sigmoidal variable slow semi-log fit with an $R^2$ = 0.9855. Each standard was measured in duplicate on each ELISA.

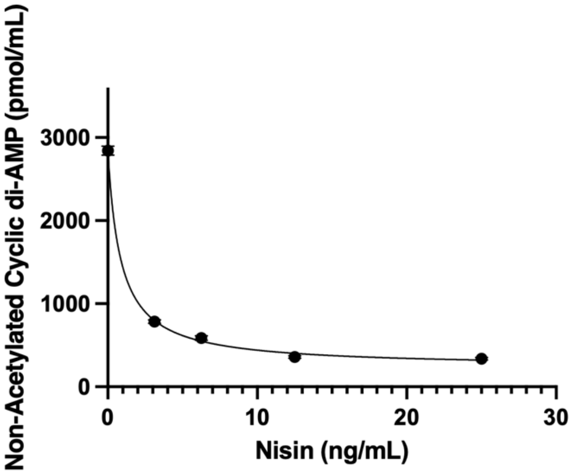


Figure S9. **EpaU induction with nisin shows a phenotype even at low protein levels**.
To correlate how much EpaU might be necessary to see an effect on c-di-AMP levels, various concentrations of nisin were used to induce EpaU in the Δ*epaU*::p*epaU* cells. Intracellular c-di-AMP concentration was measured using an ELISA. Levels of nisin far below the concentration for EpaU induction suppress c-di-AMP synthesis, suggesting that even a small amount of EpaU can reverse the phenotype. All measurements were done with three biologically independent repeats, with the line representing the median value.

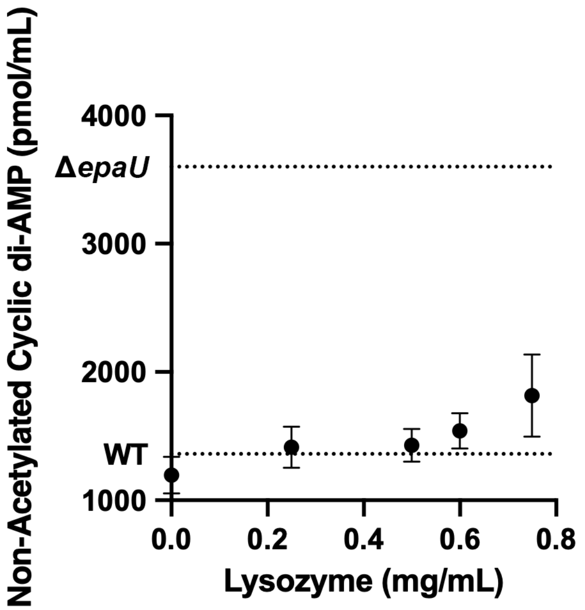


Figure S10. **Lysozyme cell wall challenge does not promote c-di-AMP level change.** Lysozyme was titrated into cultures during *E. faecalis* growth as outlined in Methods. Dashed lines mark the measured levels of WT and Δ*epaU* from Fig. 8C, respectively. Data are mean values ± S.D., n = 3 biologically independent experiments.

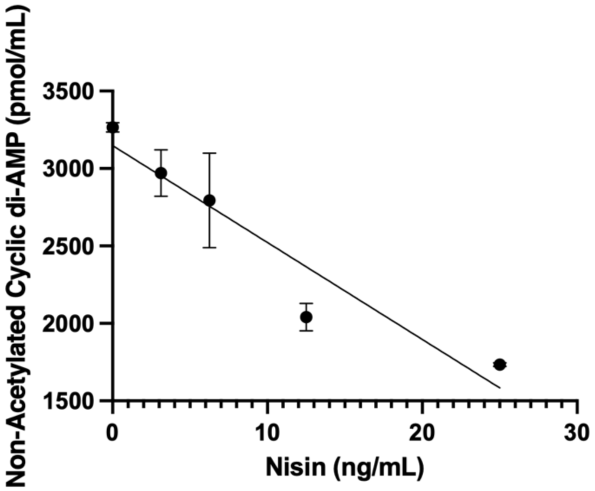


Figure S11. **Effect of expression of EpaU lacking the cell wall binding domain on c-di-AMP levels.**
A construct lacking the CWB domain, Δ*epaU*::p*epaU*ΔCWB, was induced with various levels of nisin and then assayed for intracellular c-di-AMP concentration using an ELISA. While high concentrations of nisin did show a phenotype more consistent with WT, lower levels of protein expression tend to look more like Δ*epaU*, suggesting that the CWB domain is critical for the function of EpaU at typical cellular concentrations, and activity of the catalytic only domain is only observed in the case of gross over-expression. All measurements are from three biologically independent experiments, with the trend-line representing a non-rigorous fit.

Table S1. MICs of other antibiotics tested between WT and Δ*epaU* strains.

| **MIC** | **1316 (μg/mL)** | **95% CI (μg/mL)** | **Δ*epaU* (μg/mL)** | **95% CI (μg/mL)** |
|---|---|---|---|---|
| **Ampicillin** | 6.986 | 6.390 to 7.714 | 10.61 | 9.322 to 12.38 |
| **Penicillin** | 1.833 | 1.616 to 2.104 | 1.940 | 1.672 to 2.288 |
| **Fosfomycin** | 11.39 | 10.65 to 12.34 | 12.66 | ND[a] to 17.35 |
| **Vancomycin** | 2.828 | 2.638 to 3.050 | 3.062 | 2.696 to 3.547 |
| **Tunicamycin** | 1.110 | 1.056 to 1.170 | 1.008 | ND |
| **Daptomycin** | 9.731 | 8.514 to 11.29 | 10.37 | 8.689 to 12.54 |
| **Ceftriaxone** | 13.43 | 9.783 to 21.20 | 11.82 | 8.360 to 20.73 |

[a]Values marked not defined (ND) were unable to be computed by Prism and were reported by the program as interrupted calculations.

Table S2. Best-fit values for ampicillin resistance of other control strains.

| **Best-fit values** | **WT::pMSP3535** | **WT::p*yycJ*** | **WT::p*cdaA*** | **V583** | **Δ*altA*** |
|---|---|---|---|---|---|
| logMIC | 0.7849 | 0.8246 | 0.8723 | 0.5204 | 0.5308 |
| Slope | 6.325 | 4.836 | 3.820 | 24.19 | 23.28 |
| Bottom | 0.1059 | 0.1005 | 0.09737 | 0.1158 | 0.1217 |
| Span | 0.7509 | 0.7687 | 0.7250 | 0.6858 | 0.5956 |
| MIC | 6.094 | 6.678 | 7.453 | 3.314 | 3.394 |

| **95% CI** | **WT::pMSP3535** | **WT::p*yycJ*** | **WT::p*cdaA*** | **V583** | **Δ*altA*** |
|---|---|---|---|---|---|
| logMIC | 0.7381 - 0.8365 | 0.7670 - 0.8927 | 0.7964 - 0.9670 | 0.5052 - 0.5418 | 0.5098-0.5612 |
| Slope | 5.068 - 8.091 | 3.875 - 6.150 | 3.012 - 4.939 | 17.54 - 35.48 | 15.87-37.52 |
| Bottom | 0.07909 - 0.1314 | 0.06663 - 0.1306 | 0.05418 - 0.1331 | 0.08658-0.1451 | 0.09566-0.1478 |
| Span | 0.7058 - 0.7993 | 0.7141 - 0.8289 | 0.6603 - 0.7982 | 0.6489 - 0.7231 | 0.5627-0.6290 |
| MIC | 5.471 - 6.863 | 5.847 - 7.811 | 6.258 - 9.268 | 3.200 - 3.482 | 3.234 - 3.641 |

Best-fit values for ampicillin resistance of WT::pMSP3535, WT::p*yycJ*, Δ*altA*, and WT::p*cdaA* and V583 control strains. MIC reported in μg/mL; slope, bottom, and span are parameters from Gompertz equation for MIC determination (2). Fits are the result of three independent biological repeats.

Table S3. Strains and plasmids used in this study.

| Strain or plasmid | Description [a] | Reference |
|---|---|---|
| *Enterococcus faecalis* | | |
| VE14089 | *E. faecalis* V583-derived strain cured of plasmids | (3) |
| V583 | Vancomycin-resistant clinical isolate | (4) |
| Δ*epaR* | *epaR* deletion mutant in VE14089 strain | This study |
| Δ*epaX* | *epaX* deletion mutant in VE14089 strain | This study |
| Δ*epaR*::p*epaR* | Δ*epaR* complemented with p*epaR* carrying *E. faecalis epaR*, $Cam^R$ | This study |
| Δ*epaX*::p*epaX* | Δ*epaX* complemented with p*epaX* carrying *E. faecalis epaX*, $Cam^R$ | This study |
| OG1RF | spontaneous mutant of OG1 strain, resistant to rifampicin and fusidic acid | (5) |
| YI6-1 | Clinical isolate YI6 cured of plasmid pYI6 | (6) |
| ATCC 19433 | NCTC 775 | (7) |
| X98 | fecal isolate (MLST 19) | (8) |
| Δ*cpsC* | *cpsC* deletion mutant in V583 strain | (9) |
| Δ*atlA* | *atlA* deletion mutant in V583 strain | (10) |
| Δ*epaU* | *epaU* deletion mutant in VE14089 strain | This study |
| Δ*epaU*::p*epaU* | Δ*epaU* complemented with p*epaU* carrying *E. faecalis epaU*, $Er^R$ | This study |
| Δ*epaU*::pMSP3535 | Δ*epaU* with empty complementation plasmid pMSP3535, $Er^R$ | This study |
| VE14089::p*cdaA* | WT with p*cdaA* carrying *E. faecalis cdaA*, $Er^R$ | This study |
| VE14089::p*yycJ* | WT with p*yycJ* carrying *E. faecalis yycJ*, $Er^R$ | This study |
| *Streptococcus mutans* | | |
| Δ*rgpG* | *rgpG* deletion mutant (has a nonpolar spectinomycin resistance cassette inserted in *rgpG*), $Spec^R$ | (11) |
| *Escherichia coli* | | |
| NovaBlue | *E. coli* strain for cloning | Novagen |
| MC1061 | *E. coli* strain for cloning | (12) |
| LOBSTR(DE3)pRARE2 | *E. coli* strain for protein expression; $Cam^R$ | Kerafast |
| Plasmids | | |
| pDC123 | *E. coli-streptococcus* shuttle vector, JS-3 replicon, chloramphenicol resistance cassette, $Cam^R$ | (13) |
| pLT06 | Vector for making mutant *E. faecalis* strains, $Cam^R$ | (9) |
| pMSP3535 | Vector for nisin-controlled protein expression in *E. faecalis*, $Er^R$ | (14) |
| pRSF-NT | Vector for protein expression in *E. coli* with N-terminal $His_6$ tag followed by a TEV cleavage site, $Kan^R$. | (15) |
| pKV1620 | pLT06-based vector for *epaR* deletion, $Cam^R$ | This study |
| pKV1694 | pLT06-based vector for *epaX* deletion, $Cam^R$ | This study |

| | | |
|---|---|---|
| pKV1667 | pLT06-based vector for *epaU* deletion, $Cam^R$ | This study |
| pKV1633 | p*epaR*, pDC123-based vector for Δ*epaR* complementation, $Cam^R$ | This study |
| pKV1634 | p*epaX*, pDC123-based vector for Δ*epaR* complementation, $Cam^R$ | This study |
| p*epaU* | pMSP3535-based vector for Δ*epaU* complementation, $Er^R$ | This study |
| p*epaU* E262Q | Vector with the catalytic site mutant EpaU E262Q, $Er^R$ | This study |
| p*epaU* ΔCWB | Vector with EpaU lacking the CWB domain, $Er^R$ | This study |
| p*cdaA* | pMSP3535-based vector for nisin-inducible expression of diadenylate cyclase CdaA, $Er^R$ | This study |
| p*yycJ* | pMSP3535-based vector for nisin-inducible expression of a putative metallo hydrolase YycJ (EF1197), $Er^R$ | This study |
| pKV1606 | pRSF-NT-based vector for expression of sfGFP-$EpaU^{V583}$ C-terminal domain fusion, $Kan^R$ | This study |
| pKV1608 | pRSF-NT-based vector for expression of $EpaU^{V583}$ C-terminal domain, $Kan^R$ | This study |
| pKV1695 | pRSF-NT-based vector for expression of sfGFP-$EpaU^{OG1RF}$ C-terminal domain fusion, $Kan^R$ | This study |
| pKV1696 | pRSF-NT-based vector for expression of full-length $EpaU^{V583}$ without signal peptide and N-terminal IDR, $Kan^R$ | This study |

Table S4. Primers used in this study.

| Primer | Sequence[a] | Genetic manipulations |
|---|---|---|
| epaR_F_BamH | CGCGGATCCGCGCGATTGGTGCAACGTTGATGT | construction of Δ*epaR* |
| epaR_R1_Not | ACTAGCGCGGCCGCTTGCTCCCCGTCTTGCATTACTCCATTCAC | |
| epaR_F1_Not | GGAGCAAGCGGCCGCGCTAGTGGATTGGATGAGTCTGAAGAACG | |
| epaR_R_EcoR | CCGGAATTCCGGTCCACTTTTCCTTTTCCTCCCT | |
| epaR_check_F | GAAACCGCTGGGGACACATA | verification of Δ*epaR* |
| epaR_check_R | TCTGGTTGAATTTCAAAATTTGAGACT | |
| 2170_BamH_F | GTAGGATCCATGCGAATATTTCGATCTGGG | construction of Δ*epaX* |
| 2170_armR | TAAAATCATTGTTTTTCTAAATATTTTTCTACATTGTATACAGGAAC | |
| 2170_armF | AGAAAAATATTTAGAAAAACAATGATTTTACTTATGTTTGGTCTTCC | |
| 2170_EcoR_R | GTTGAATTCCGAAAACAATAGCACCACAG | |
| 2170_chF | GAGAAGCCACCGTCCAAG | verification of Δ*epaX* |
| 2170_chR | CACCTGACAATAGCATAATTATCAAGG | |
| 2174_BamH_F | GAGAGGATCCAGATTATGAAAAATTGAAAGTGC | construction of Δ*epaU* |
| 2174_R2 | TATTAGCACTAAAACTATTTAATAATACAGTACAAATTAACATACC | |
| 2174_F2 | ATTATTAAATAGTTTTAGTGCTAATAAGTGGTATGTAGCAAAAGTG | |
| 2174_EcoR_R | GAATGAATTCGAATGAGCTGCTACAGGATATTAAC | |
| 2174_checkF | TGAAATAAAGGTATGAGATAATAATTGGTG | verification of Δ*epaU* |
| 2174_checkR | AAAAACACCTTTGTCGAACTCG | |
| 2177_Xba | CTATCTAGAATTACTGGAAGTGAAGAGAGGAAG | construction of p*epaR* |
| 2177_BamH | CTTGGATCCTTAACGGTAAATTCTAATAGCATTCTCTTTAC | |
| 2170_Xba | CTATCTAGAATTTTAGAAGGTAGCGGAAATTGATATG | construction of p*epaX* |
| 2170_BamH | CTTGGATCCTTAATCTATTGTTTTTTCTATATATTTTCTTTTCAATTTAATTGG | |
| epaU-BamH-F | CCAGGATCCATATTATTAAATATATAAGGAGAGTTTCATG | construction of p*epaU* |
| epaU_Xho | CTTCTCGAGCTTAACCCACTTTTGCTAC | |
| epaU_E262Q_F | TAATGATGCGCAAGATCCAACGCTTACAAAC | construction of p*epaU* E262Q |
| epaU_E262Q_R | GTTGGATCTTGCGCATCATTAAACATTATGG | |
| epaU_FLAG_F | CGGTGATTATAAAGATCATGATATCGATTACAAGGATGACGATGACAAGTAAGGGCCTGACTTAAGTAAG | construction of p*epaU* ΔCBD |
| epaU_FLAG_R | ATATCATGATCTTTATAATCACCGTCATGGTCTTTGTAGTCACTACTATTACCCATATTTAAAAAATTAC | |
| cdaA_BamH | GATGGATCCGGTATACTTGTTAGGAAATCTTGAG | construction of p*cdaA* |
| cdaA_Xho | ATCCTCGAGTCATTTGCGTTTAACCCCC | |
| mbl_Bgl2 | GATTAGATCTGAAACACGTTAATGTTGTAATGTAG | construction of p*yycJ* |
| mbl_Xho | ATCCTCGAGCTAAATTTGAAATAACTCTGAGGC | |
| sfGFP_Nco | GAGACCATGGCTAGCAAAGGAGAAGAACTTTTC | |

| | | |
|---|---|---|
| gfp_2174_R | CTTACTTAAGTCAGGCCCACTACCTCCACCTTTGTAGAGC | construction of pKV1606 |
| gfp_2174_F | GCTCTACAAAGGTGGAGGTAGTGGGCCTGACTTAAGTAAG | |
| 2174_Hind | CTCAAGCTTAACCCACTTTTGCTACATAC | |
| 2174_374Nco | GAGACCATGGGGCCTGACTTAAGTAAGTATTAC | construction of pKV1608 |
| 2174_Hind | CTCAAGCTTAACCCACTTTTGCTACATAC | |
| sfGFP_Nco | GAGACCATGGCTAGCAAAGGAGAAGAACTTTTC | construction of pKV1695 |
| U_og1rf-R1 | GGAATTGCCACCTCCACCTTTGTAGAGC | |
| U_og1rf-F1 | GCTCTACAAAGGTGGAGGTG | |
| U_og1rf-R | GGTGCTCGAGTGCGGCCGCAAGCTTAATTC | |
| 2174_113BspH | GAGATCATGACCATTGACCAAGATGAAGC | construction of pKV1696 |
| 2174_Hind | CTCAAGCTTAACCCACTTTTGCTACATAC | |

[a] Restriction sites are underlined.
